# Impact of virus subtype and host *IFNL4* genotype on large-scale RNA structure formation in the genome of hepatitis C virus

**DOI:** 10.1101/2020.06.16.155150

**Authors:** P. Simmonds, L. Cuypers, W.L. Irving, J. McLauchlan, G.S. Cooke, E. Barnes, STOP-HCV Consortium, M.A. Ansari

## Abstract

Mechanisms underlying the ability of hepatitis C virus (HCV) to establish persistent infections and induce progressive liver disease remain poorly understood. HCV is one of several positive-stranded RNA viruses capable of establishing persistence in their immunocompetent vertebrate hosts, an attribute associated with formation of large scale RNA structure in their genomic RNA. We developed novel methods to analyse and visualise genome-scale ordered RNA structure (GORS) predicted from the increasingly large datasets of complete genome sequences of HCV. Structurally conserved RNA secondary structure in coding regions of HCV localised exclusively to polyprotein ends (core, NS5B). Coding regions elsewhere were also intensely structured based on elevated minimum folding energy difference (MFED) values, but the actual stem-loop elements involved in genome folding were structurally entirely distinct, even between subtypes 1a and 1b. Dynamic remodelling was further evident from comparison of HCV strains in different host genetic background. Significantly higher MFED values, greater suppression of UpA dinucleotide frequencies and restricted diversification were found in subjects with the TT genotype of the rs12979860 SNP in the *IFNL4* gene compared to the CC (non-expressing) allele. These structural and compositional associations with expression of interferon-λ4 were recapitulated on a larger scale by higher MFED values and greater UpA suppression of genotype 1 compared to genotype 3a, associated with previously reported HCV genotype-associated differences in hepatic interferon-stimulated gene induction. Associations between innate cellular responses with HCV structure and further evolutionary constraints represents an important new element in RNA virus evolution and the adaptive interplay between virus and host.

## INTRODUCTION

Hepatitis C virus (HCV) is a major pathogen of humans infecting more than 71 million individuals. An estimated 400,000 deaths per year occur as a direct consequence of progressive inflammatory disease and hepatocellular carcinoma over many decades of persistent infection (Collaborators, 2017; Westbrook and Dusheiko, 2014). HCV is one of a small number of RNA viruses capable of establishing long term infections in humans and other vertebrates (Hoofnagle, 2002). This contrasts with the more typical outcomes of RNA virus infections - acute infection and rapid clearance mediated through the actions of innate and adaptive immune responses and subsequent protective immunity. How HCV is able to counteract the otherwise powerful host responses mounted against viruses has remained unclear for several decades since the concept of RNA virus persistence became recognised (Oldstone, 2009; Randall and Griffin, 2017).

Persistence is the observed outcome of infections with several other mammalian viruses including human pegivirus (HPgV) and related viruses in the *Hepacivirus* and *Pegivirus* genera of *Flaviviridae* infecting other mammals (Baechlein *et al.*, 2015; Pfaender *et al.*, 2015; Ploss *et al.*, 2009; Trivedi *et al.*, 2017). It is also documented in several picornaviruses (*eg.* foot-and-mouth disease virus; FMDV; (Condy *et al.*, 1985; Cortey *et al.*, 2019)) and caliciviruses (murine norovirus [MNV] and some vesiviruses) (Coyne *et al.*, 2006; Forrester *et al.*, 2003; Hsu *et al.*, 2006; Thackray *et al.*, 2007). Collectively, they differ substantially in many aspects of their host interactions and disease outcomes. Infections with several are asymptomatic, as in the case of HPgV and, where known, in other animal pegiviruses and in MNV. While FMDV may cause severe and often fatal vesicular disease in cows and other ruminants, infections in buffalo, its natural host, is often clinically inapparent (Thomson *et al.*, 1992). Chronic and progressive liver disease is observed in infections with HCV and with some hepaciviruses infecting other host species, such as GBV-B in tamarins (Beames *et al.*, 2001; Bukh *et al.*, 2001) and equine hepaciviruses in horses (Pfaender *et al.*, 2015; Ramsay *et al.*, 2015). However, bovine hepacivirus infections of cows have been reported as entirely apathogenic (Baechlein *et al.*, 2019).

While these features of infection are disparate in both pathogenicity and target tissues, a unifying characteristic of these persistent RNA viruses is their intensely structured RNA genomes, where a high degree of internal sequence complementarity creates elaborate tandem arrays of stem-loops and potentially tertiary structure elements spanning most of the genomic RNA (Davis *et al.*, 2008; McFadden *et al.*, 2013; Simmonds *et al.*, 2008; Simmonds *et al.*, 2004). Existing bioinformatic and physicochemical analysis of this genome attribute, that we termed genome-scale ordered RNA structure (GORS) reveals many differences from the better characterised discrete elements of folded RNA found in RNA virus genomes. The latter may serve as replication elements or mediate ribosomal interactions in translation initiation or control (*eg.* frame shifting). Contrastingly, the distributed and extensive nature of GORS leads to a major difference in the configuration of the RNA in terms of its overall shape and accessibility to hybridisation to external probes (Davis *et al.*, 2008). How this globally folded configuration of genomic RNA contributes to the interaction of the virus and its host (and its ability to establish persistence) remains unknown.

In the current study we have re-investigated many aspects of GORS in HCV infections now that there are many thousands of accurately determined whole genome sequences of HCV available. Furthermore, RNA structure information on pairing interactions is now available for several whole genome RNA molecules using recently developed SHAPE methods (Mauger *et al.*, 2015; Pirakitikulr *et al.*, 2016). In the current study, we have analysed the nature and phylogenetic conservation of RNA structures in HCV and their commonality in extents and constraints with RNA folding in the genomes of other persistent viruses. We show that the formation of RNA structure may be a much more dynamic evolutionary process that may furthermore evolve in response to genetic background of the host it infects.

## RESULTS

### RNA structure prediction in HCV genomes

The propensity of RNA to internally base pair and its structural configuration is dependent on both the order of bases and the G+C content of the sequence. The sequence order component of RNA structure formation in HCV genomes was estimated by comparison of minimum folding energies (MFEs) of native sequences with those of the same sequence scrambled in base order while maintaining native dinucleotide frequencies using the algorithm NDR. The resulting values (MFEDs) were generated for consecutive 240 base fragments incrementing by 9 bases through the coding region of whole genome sequences of HCV genotypes 1a (n=388), 1b (n=106) and 3a (n=855). MFED values greater than zero were observed throughout the genomes of each genotype analysed (Fig. 1A), although there were some differences in the degree of MFED elevation predicted for different genome regions. These differences were mirrored in a parallel plot of Z-scores representing the MFE of the native sequence in the distribution of MFEs of the sequence order randomised controls (Fig. S1A; Suppl. data).

**FIGURE 1.**
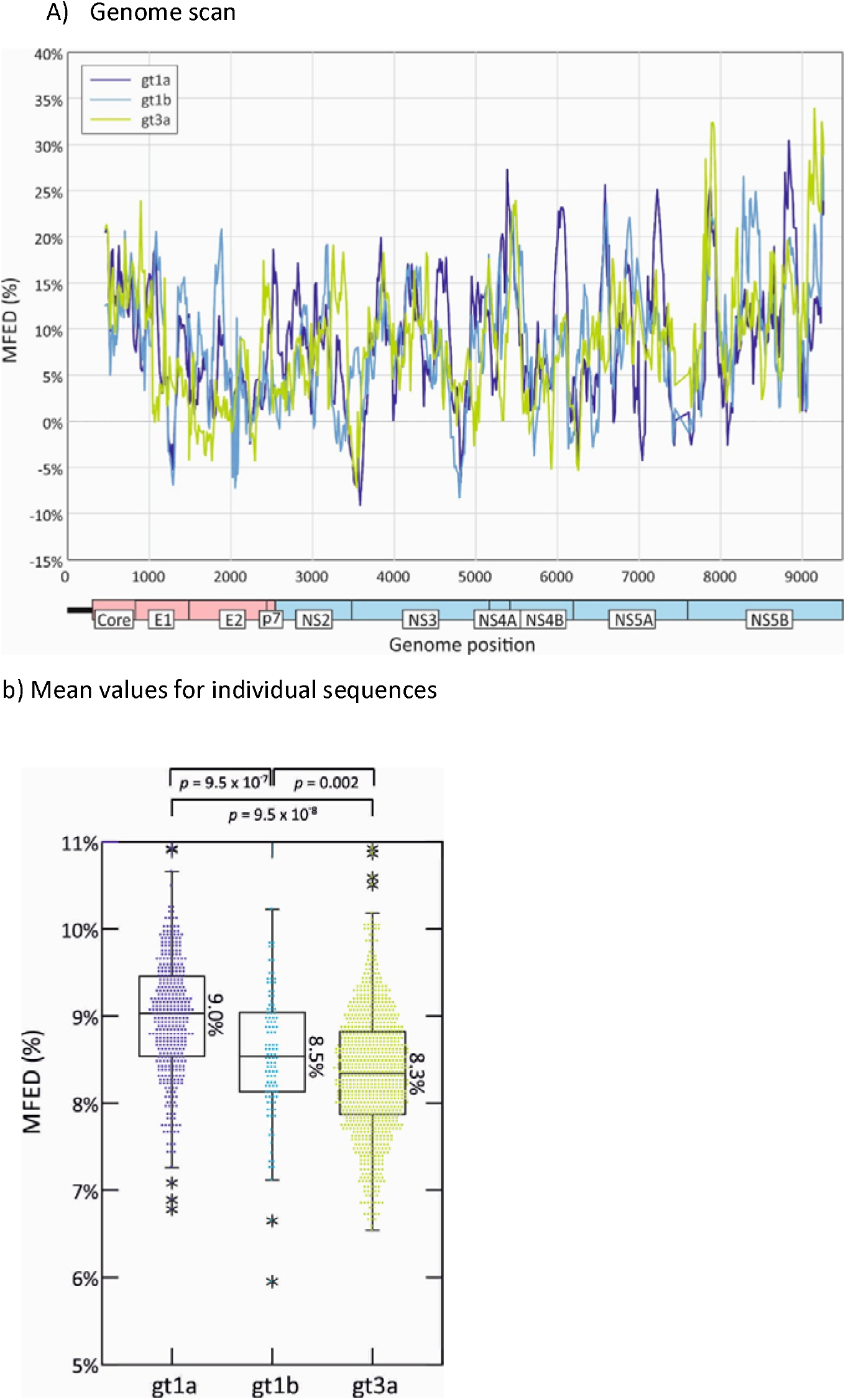
MFED DIFFERENCES IN HCV CODING REGIONS OF GENOTYPES 1a, 1a AND 3a. MFED values calculated for consecutive 240 base fragments, incrementing by 15 bases across the HCV coding region (569 fragments). MFED values were calculated by subtraction of the MFE of the native sequence by the mean value of 49 sequence order randomised controls (A) Mean values of genotypes 1a (n=388), 1b (n=106) and 3a (n=855) sequences plotted by genome position. (B) Mean values of sequence fragments for individual polyprotein sequences of gts 1a, 1b and 3a. The box plots show (from the top, 2 standard deviations (SDs) above mean, 1 SD above mean, mean, 1 SD below mean and 2 SDs below mean; stars represent outliers outside this range. Genome positions were normalised by reference to the H77 prototype sequence, AF011751 (coding region positions 342-9377). Distributions of MFED values of different genotypes were compared by Kruskall Wallace non-parametric test; *p* values on comparing 1a with 1b, 1a with 3a and 1b with 3a are shown above graph.

Overall, however, mean MFED values for whole coding regions sequences were observed in sequences from all genotypes (6.0%-10.9% range in MFED values) and more strikingly, between variants of the same genotype (Fig. 1B). Although there was a scatter of MFED values within each genotype, MFED values were significantly higher in genotype 1a than 1b and 3a (Fig. 1B). Genotype-associated differences in predicted RNA structure formed by the three genotypes were observed across the genome; MFED values calculated from individual fragments of gt1a and gt3a sequences frequently differed from each other throughout the coding region (Fig. S1B; Suppl. Data); they were not confined to individual stem-loops. Increased MFED values were also observed irrespective of the underlying degree of sequence diversity of HCV in different parts of the genome. For example, they remained elevated throughout the E1/E2 region despite the substantially greater sequence variability of the envelope protein gene sequences, particularly around the hypervariable region of E2 (Fig. S1C; Suppl. Data).

The use of mean values of bulk MFED values for collections of sequences of the same subtype or genotype provides a very coarse-grained indication of the RNA structural variability of HCV at the genotype or individual sequence level. To visualise structural heterogeneity more effectively, we developed a new plotting method for RNA structures predicted by RNAFOLD. These were based on sampling an ensemble of sub-optimal folds generated by the program SubOpt.exe using predictions for individual sequences and recording pairing predictions supported by 50% or more of the ensemble. For calculating stem-loop heights, unpaired bases in terminal loops of each secondary structure were identified in the ensemble consensus connect file for each sequence and plotted as a height of zero (z-axis) with their genome positions and sequence number in x- and y-axes. Neighbouring bases were successively plotted according to a colour scale that reflects their distance in the stem from the terminal loop. Examples of a standard stem-loop, an interrupted stem-loop and a more complex clover leaf structure are shown in their MFOLD representation and 3 and 2-dimensional contour plots (Fig. 2A, 2B and 3C respectively). Contour Plots therefore provide an approximate visualisation of the positions, shapes and sizes of RNA structure elements across whole alignments of potentially large numbers of sequences. Applying this method to a short section of the HCV genome in the core / E1 gene region (Fig. 2D), the 3-dimensional displays sequence positions (x-axis), sequence number in the alignment (y-axis) and relative heights of predicted stem-loops. Its transformation to a 2-dimensional representation and associated depiction of structural heterogeneity between sequences greatly facilitates an analysis of RNA structure conservation between and within genotypes.

**FIGURE 2.**
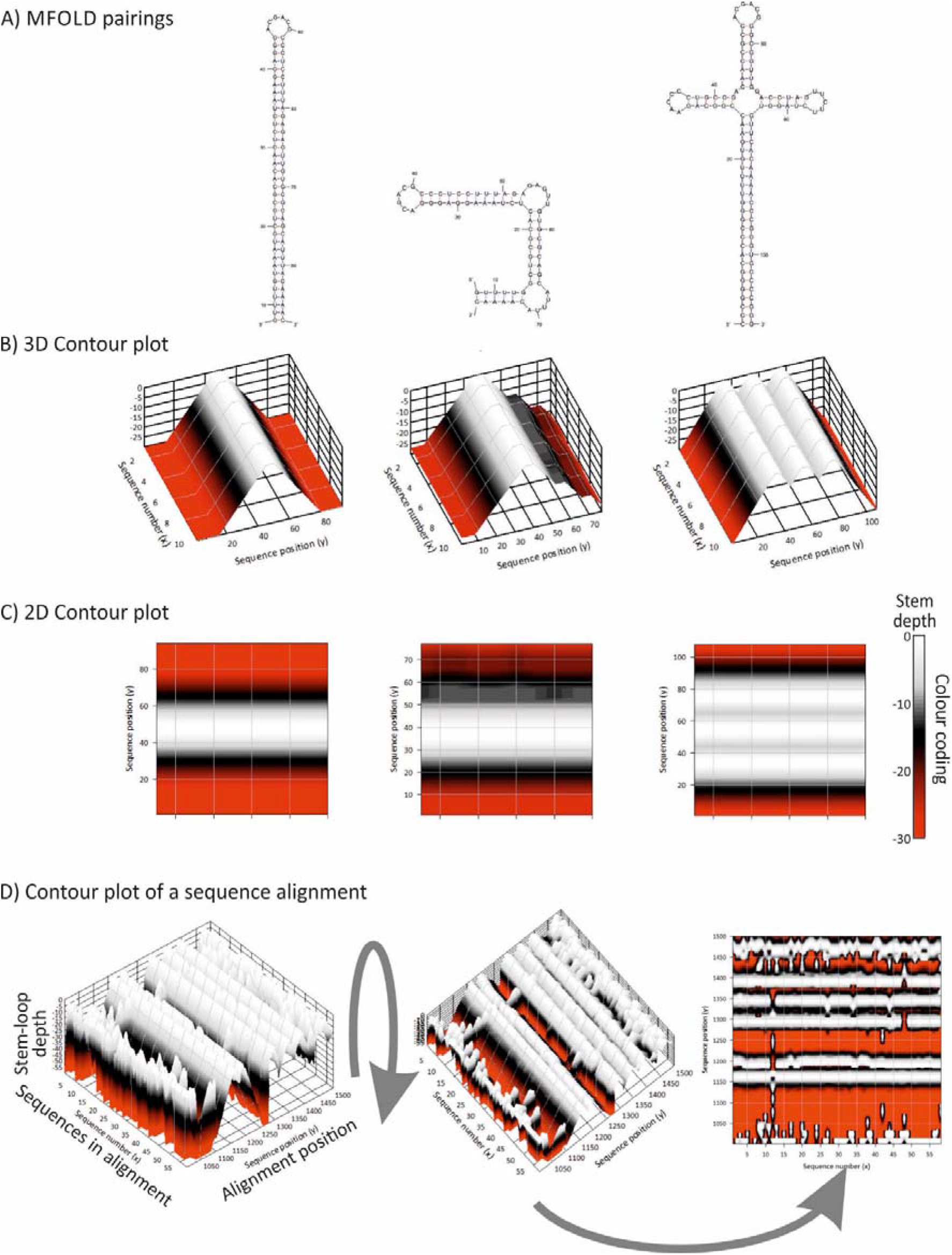
VISUALISATION OF RNA STRUCTURE CONSERVATION IN A SEQUENCE ALIGNMENT. Visualisation of RNA structure conservation in a sequence alignment. Comparison of RNA structure visualisation using (A) MFOLD, (B) 3D and (C) 2D contour plots produced by StructureDist of example RNA secondary structure elements (stem-loop, interrupted stem-loop and clover leaf). Predicted consensus positions of terminal loops in RNAFold RNAsubopt output were aligned and plotting depths based on pairings either side calculated.. There were depicted as canyons corresponding to duplex lengths (B) and then rotated to create a 2D representation of sequences in the alignment (x-axis) and alignment position (y-axis) and colour-coded depth. (D) Application of contour plotting to an alignment of HCV sequences in the core / E1 region in 3 and 2-dimensional representations. Pairing predictions represent as majority role consensus derived from the ensemble of sub-optimal folds produced by the RNAsubopt program; similar results are obtained irrespective of whether suboptimal folds are sampled from a defined range of MFE values from the optimum, suboptimal structures sampled based on their Boltzmann weights in a partition function or Zuker suboptimals (Fig. S2B; Suppl. Data).

Much larger scale contour plots of the coding region sequences of samples obtained in the current study from gt1a, gt1b, gt2a and gt3a (supplemented with published sequences for gt2a) were created from 13 sequential 1600 base fragments of each alignment incrementing by 400 bases between fragments. The fragment size of 1600 bases chosen for individual contour plots reproduces pairings predicted from shorter and longer sequence fragments (Fig. S2A; Suppl. Data); this represents a necessary compromise between computational time and enabling the method to detect longer range-base pairings. Using these settings, this composite genome-wide representation revealed the existence of highly conserved structure elements across genotypes located specifically in the core and NS5B regions (Fig. 3A, 3B). These were highly concordant with RNA pairings detected by SHAPE mapping for genotypes 1a, 1b and 2a in a previous study (Mauger *et al.*, 2015) replotted in the same contour format (Fig. 3). RNA structure determination by SHAPE (Mauger *et al.*, 2015; Pirakitikulr *et al.*, 2016) indeed largely verifies previously described RNA structure prediction programs and analysis of co-variance (Tuplin *et al.*, 2004) and RNase mapping (Tuplin *et al.*, 2004). Several of the RNA secondary structures have been functionally characterised through investigation of effects on replication when disrupted (Diviney *et al.*, 2008; Mauger *et al.*, 2015; McMullan *et al.*, 2007; Pirakitikulr *et al.*, 2016; You *et al.*, 2004) or through measurement of replication effects of systematic large scale sequence mutation in different coding region segments (Chu *et al.*, 2013). The simplified summary in Fig. 3A highlights the concentration of investigated structures in the genome ends. It also demonstrates the relative infrequency with which identified structures influence replication capacity in cell culture.

**FIGURE 3.**
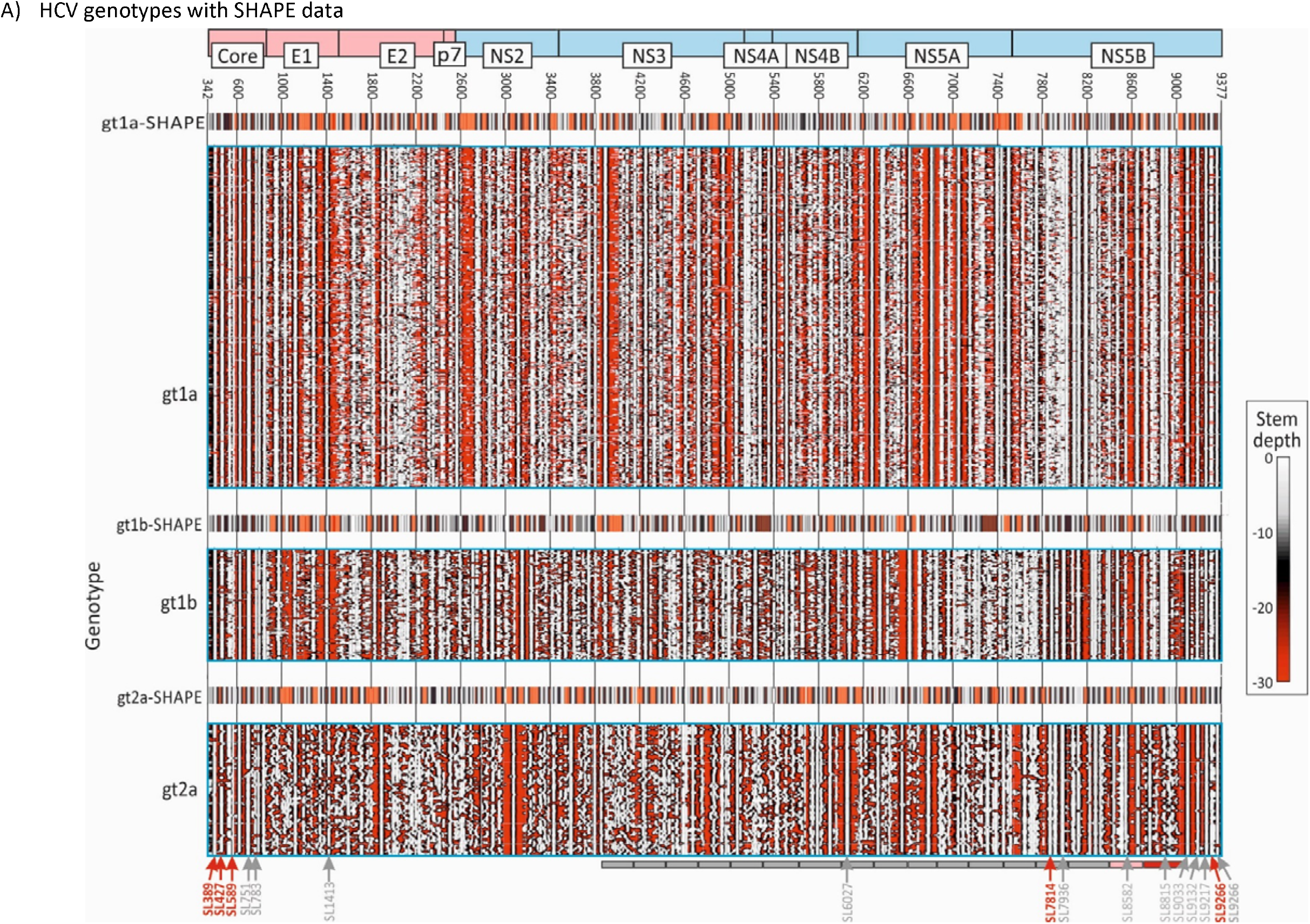
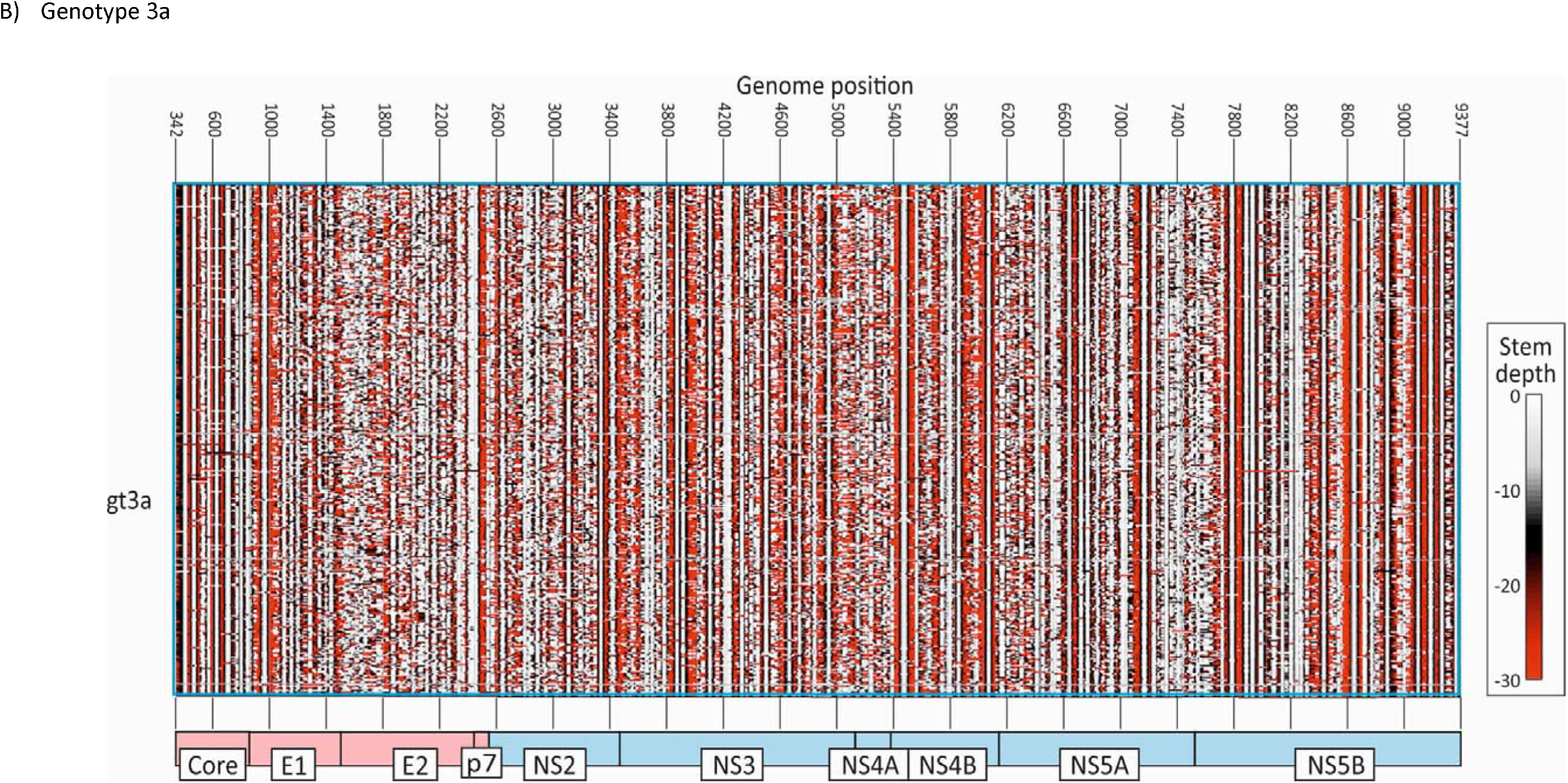
COMPARISON OF RNA STRUCTURE PREDICTIONS FOR HCV GENOTYPE 1a, 1b, 2A AND 3a SEQUENCES SHOWN AS CONTOUR PLOTS. Contour plots of RNA structure predictions by RNAFOLD of the coding regions of (A) 388 gt1a, 106 gt1b, 66 gt2a and (B) 855 gt3a sequences obtained in the current study. Inset plots show RNA structures previously determined for gt1a (H77), gt1b (Con1) and gt2a (JFH1) RNA transcripts by SHAPE (Mauger *et al.*, 2015) plotted to the same scale. The contour plot scale and genome positions of RNA structure elements were converted to those in the H77 (gt1a) prototype sequence (AF011751). The positions of functionally investigated RNA structures of genome segments in previous studies (#27362; #26127; #28018; #31034; #31045; #31046} are indicated by arrows underneath the contour plot of gt2a; numbering and arrows point to the first (5’) bases of the stem-loop. Those show little or no phenotypic effect on disruption shown in grey; those leading to substantial (>1 log) reductions in replication shown in red. Phenotypes of viruses with 17 systematically mutated segments across the NS region (Chu *et al.*, 2013) indicated in grey (no effect), grey (mild effect – segment 16) and red (severe effect - segment 17). These functional depiction is a simplification and interactions between RNA structures with other elements in coding and non-coding regions have been excluded; for this information please consult the original publications.

However, the most striking aspect of the structure predictions is the extensive variability in the position and intensity of structure formation between genotypes and subtypes of HCV. In regions outside of the core and 3’terminal NS5B genes, structure conservation was evident at subtype level only, with only a small number of predicted stem-loops shared across two or more subtypes. There were additionally regions that showed extensive predicted structural heterogeneity within subtypes, including the envelope genes and parts of NS5A and NS5B. Despite these structural differences, areas without sequence conservation between subtypes nevertheless showed elevated MFED values (Fig. 1A), indicating that in the period in which the 4 subtypes had diversified, there has been a substantial degree of RNA structure re-invention and appearance of quite different structured elements while at the same time, presumably maintaining similar levels of overall structure (Fig. 1B). Further structural diversity of HCV is apparent on examination of representative examples of each currently classified HCV subtype (Fig. S3; Suppl. Data), where the only shared structural elements are the stem-loops in the core and 3’ NS5B regions. Despite the sequence divergence (>30%) and structural heterogeneity evident from the contour plots, each currently classified HCV subtype showed elevated MFED values, but with some variability between genotypes in mean values and ranges (Fig. S4; Suppl. Data).

The ability of contour plots to localise areas of RNA secondary structure was investigated by extension of the analysis to virus groups with previously documented structured genomes and for which full genome sequences from a range of strains has been previously obtained (foot-and-month disease virus [FMDV] type O, human pegivirus type 1 [HPgV-1] and murine norovirus type 3 [MNV3] analysed in the current study) (Davis *et al.*, 2008; McFadden *et al.*, 2013; Simmonds *et al.*, 2008; Simmonds *et al.*, 2004)(Table 1). Contour plots of these were compared with example datasets of unstructured virus groups (enterovirus A71 [EV-A71], human parechovirus type 3 [HPeV-3] and Japanese encephalitis virus [JEV]) (Fig. 4). These virus groups were selected to possess similar degrees of naturally occurring sequence divergence as found within the gt1a, 1b, 2a and 3a datasets (Table 1) which might otherwise influence degrees of RNA structure conservation. Similarly to HCV, the structured viruses (FMDV-O, HPgV-1 and MNV-3) showed evidence for conserved areas of RNA structure formation throughout their genomes, with substantial ordering of stem-loop structures in large parts of the genome. HPgV-1, which possesses the highest MFED values, shows a series of very large and quite regularly spaced stem-loops throughout the entire coding region. FMDV similarly shows packed RNA structures although with shorter stem-loops than observed in HPgV-1. MNV has a lower overall MFED value than the other structured viruses and possesses fewer regions of ordered structure than FMDV and HPgV-1. It is also apparent that some conserved loops (*eg.* at positions 5045) may play functional roles as elements of the sub-genomic promoters for the capsid gene (Simmonds *et al.*, 2008). Further structures may contribute to the expression of ORF3 genes and frameshifting for ORF4. As with HCV, extension of structure prediction to a wider range of variants of FMDV variants and types introduced substantially greater heterogeneity into the structure predictions, with marked differences in structure predictions between FMDV serotypes A, C and O in the more divergent structural genes VP4, VP2, VP3 and VP1 (Fig. S5; Suppl. Data), despite there being elevated MFED values throughout this region.

**TABLE 1.** SEQUENCE DATASETS USED IN THE STUDY

| **Virus group** | **n** | **MFED** | **Het.score<sup>1</sup>** | **MPD<sup>2</sup>** |
| --- | --- | --- | --- | --- |
| HCV gt1a | 388 | 8.96% | 6.33 | 8.25% |
| HCV gt1b | 106 | 8.53% | 6.32 | 9.38% |
| HCV gt2a | 67 | 7.70% | 7.42 | 10.91% |
| HCV gt3a | 855 | 8.35% | 6.00 | 8.51% |
| EV-A71 | 138 | 0.68% | 9.75 | 14.77% |
| HPeV-3 | 38 | 1.62% | 9.12 | 9.17% |
| JEV | 65 | 1.40% | 8.72 | 8.76% |
| FMDV-O | 71 | 11.71% | 5.75 | 10.60% |
| MNV | 63 | 7.09% | 7.59 | 10.19% |
| HPgV-1 | 100 | 11.99% | 4.93 | 12.24% |

<sup>1</sup>Mean difference in predicted pairing depth between sequences at each site within a dataset; serves as a metric of heterogeneity of pairing predictions between sequences.
<sup>2</sup>Mean pairwise nucleotide sequence p distances (all sites) between members of each sequence group.

**FIG. 4.**
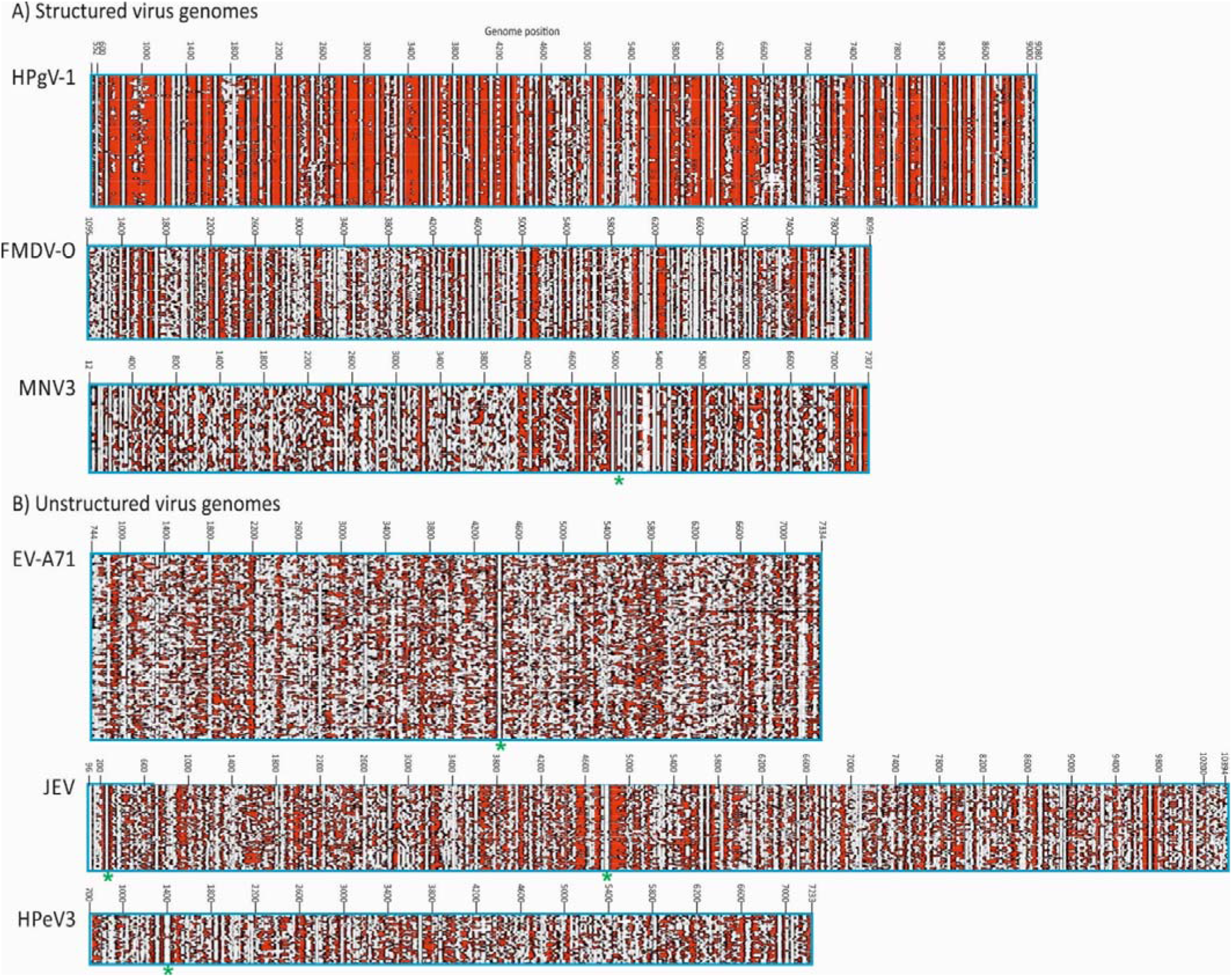
CONTOUR PLOTS OF OTHER STRUCTURED & UNSTRUCTURED VIRUS GENOMES. Comparison of contour plots of (A) structured (MFED > 5%) and (B) unstructured (MFED < 2%) virus groups plotted to scale. These show similar levels of sequence diversity (8%-15%; Table 1). Previously documented structures indicated by asterisks – EV-A71: cis-replication element (Goodfellow *et al.*, 2003; Rieder *et al.*, 2000); JEV: core region stem-loops; MNV: sub-genomic promoter (Simmonds *et al.*, 2008).

The contour plots for HCV and the three structured virus datasets (MFED values 7.7%-12.0%) were quite distinct from those of unstructured viruses (MFED values ranging from 0.7-1.6%) (Table 1). For EV-A71, large parts of the coding region showed no conserved structure formation, the exception being the *cis*-replicating element at position 4442 (Goodfellow *et al.*, 2003; Rieder *et al.*, 2000). The contour plot for JEV is similarly largely unstructured but core loops associated with translation and a large structure at position 4400 (no documented function) are evident. The HPeV-3 CRE (Al Sunaidi *et al.*, 2007) is similarly evident at position 1400 but the genome possesses few other conserved stem-loops.

The contour plot program was also used to quantify the degree of heterogeneity in folding between different sequences in an alignment. The degree of folding heterogeneity can be calculated as the mean difference in folding depth on pairwise comparisons of sequences in an alignment. Sequences in areas of conserved pairings show the same folding depths and a calculated heterogeneity of zero; pairing depths in unstructured RNA are arbitrary and therefore heterogeneous. Mean values for the whole alignments of unstructured viruses were around 9 while structured viruses showed degrees of conservation related to their MFED values (Table 1; Fig. 5). As a control, sequences of HCV genotype 1a were permuted by the algorithm CDLR to scramble codon order while retaining amino acid sequence of the encoded polyprotein and preserving native dinucleotide frequencies. It did, however, largely disrupt RNA secondary structures within the mutated region, with a reduction in mean MFED value from 9.0% to 0.5%. This change was paralleled by a large increase in heterogeneity in pairing predictions (Fig. 5). Contrastingly, little change in MFED values or pairing heterogeneity was observed on CDLR scrambled EV-A71 sequences, verifying the virtual absence of RNA secondary structure in the native sequences of this virus. The agreement between these two distinct metrics of RNA folding - sequence order-dependence and heterogeneity of predicted pairings (R^2^ = 0.93) supports the value of MFED values as a bulk metric of RNA structure formation in viral sequences.

**FIGURE 5.**
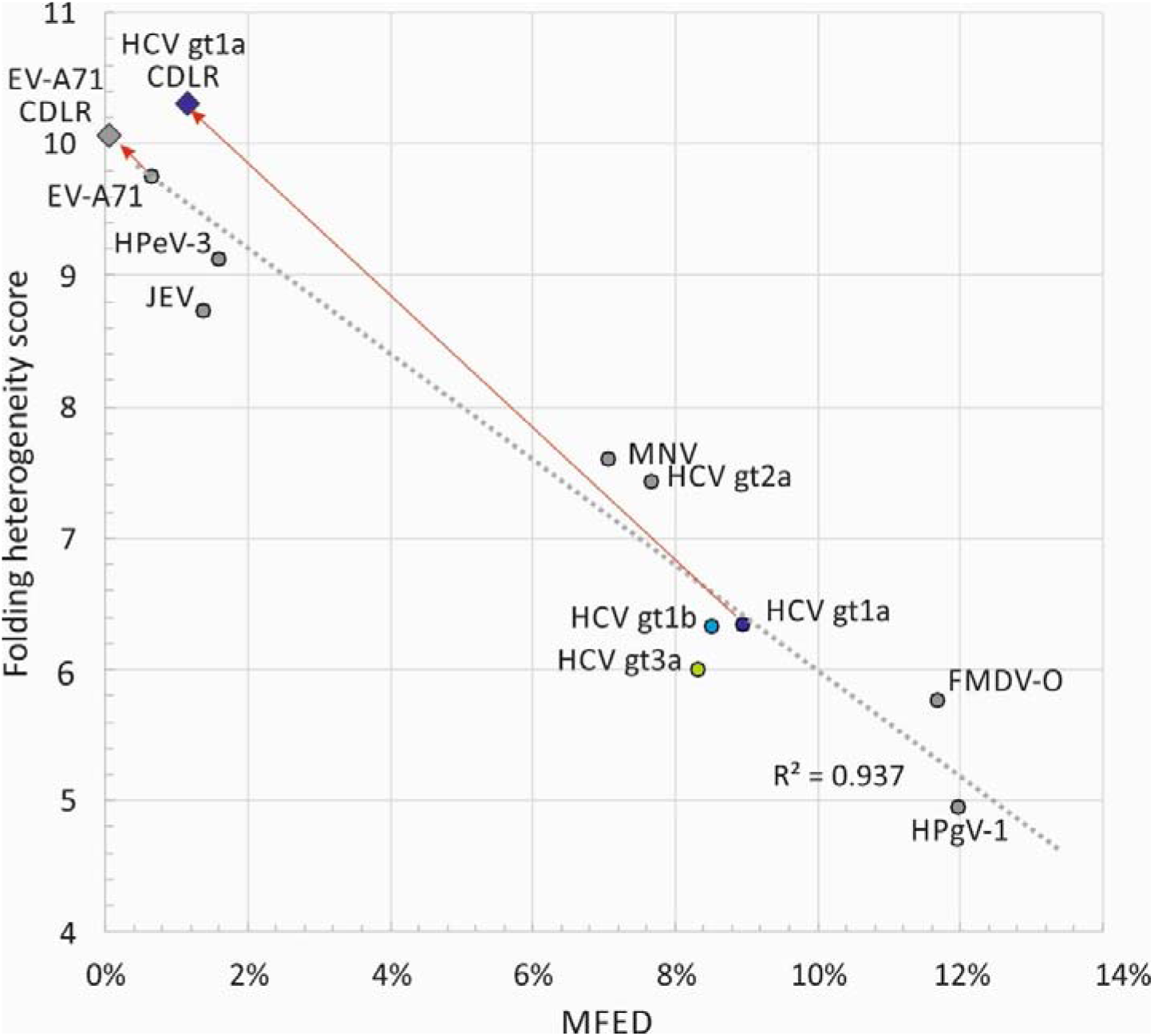
ASSOCIATION OF MFED SCORES WITH FOLDING PREDICTION HETEROGENEITY. Comparison of MFED scores with folding prediction heterogeneity for complete polyprotein sequences of HCV, other predicted structured viruses (HPgV-1, FMDV-O and MNV) and unstructured viruses (JEV, HPeV-3 and EV-A71). Folding heterogeneity was expressed as the difference in folding depth between different sequences within an alignment averaged over the whole polyprotein sequence. As controls, sequence order randomised sequence datasets were generated by the algorithm CDLR for structured (HCV-1a) and unstructured (EV-A71) genome sequences and their folding energies (MFED score; x-axis) and structural heterogeneity (y-axis) re-estimated.

### Influence of host on HCV RNA structure formation

Although different HCV genotypes and subtypes showed distinct distributions of MFED values (Fig. 1B; Fig. S4, Suppl. Data), there remained a substantial variability and overlap in MFED values between categories. To investigate whether host factors also influenced RNA structure formation, the relationship between several host demographics and clinical features with MFED values were investigated by multivariate analysis. This was based upon available data from genotype 3a individuals (n=503) in the BOSON cohort (Table 2). We additionally used information on naturally occurring differences in interferon λ4 expression from the IFNL4 gene inferred from the single nucleotide polymorphism (SNP) rs12979860, where CC, CT and TT alleles associated with no expression, medium and high-level expression respectively (Thomas *et al.*, 2009). The SNP was included in the analysis because of its previously reported effects on disease progression, outcomes of treatment and viral loads (Suppiah *et al.*, 2009; Tanaka *et al.*, 2009; Thomas *et al.*, 2009) and associations with potential drivers of sequence diversification in this dataset (Ansari *et al.*, 2019).

**TABLE 2.** ANALYSIS OF HOST FACTORS INFLUENCING MFED VALUES BY MULTIVARIATE ANALYSIS^1^

| Variable | Values/range | F-ratio | p value |
| --- | --- | --- | --- |
| IFNL4 genotype | CC, CT, TT | 9.688 | **8 X 10<sup>-5</sup>** |
| SVR | Yes, No | 1.1 | 0.295 |
| Race | White, Black, Asian<sup>2</sup> | 0.51 | 0.801 |
| Cirrhosis | Yes, No | 0.315 | 0.575 |
| Gender | Male, Female | 0.075 | 0.784 |
| Age | 19 – 70 years | 4.036 | **0.045** |
| Log VL | 10<sup>3.9</sup> - 10<sup>7.6</sup> RNA/ml | 8.7 | **0.003** |

<sup>1</sup>Analysed by the General Linear Model program in the SYSTAT v13 package
<sup>2</sup>Additional categories were indigenous North American and Indigenous Hawaiian and unknown

Multivariate analysis indeed demonstrated that IFNL4 SNP rs12979860 was the strongest predictor of MFED values, followed by viral load but no other significant host predictive factors other than a minor effects of patient age. The combination of factors accounted for approximately 25% of the variability in MFED values (multiple R = 0.267). Mean MFED values were significantly different between host *IFNL4* genotypes in both HCV genotypes 1 and 3 (Fig. 6). The direction and effect sizes were also consistent with approximately a 0.4% increase in MFED values from CC to TT genotypes in both HCV genotypes. *IFNL4* genotypes also showed substantial effects on viral diversity and UpA dinucleotide frequencies (Figs. 7, 8). There were generally highly significant differences between subjects with CC, CT and TT alleles in the sequence divergence of their infecting strains from a reconstructed ancestral sequence of each genotype, both at non-synonymous sites and synonymous sites for all three genotypes (Fig. 7).

**FIGURE 6.**
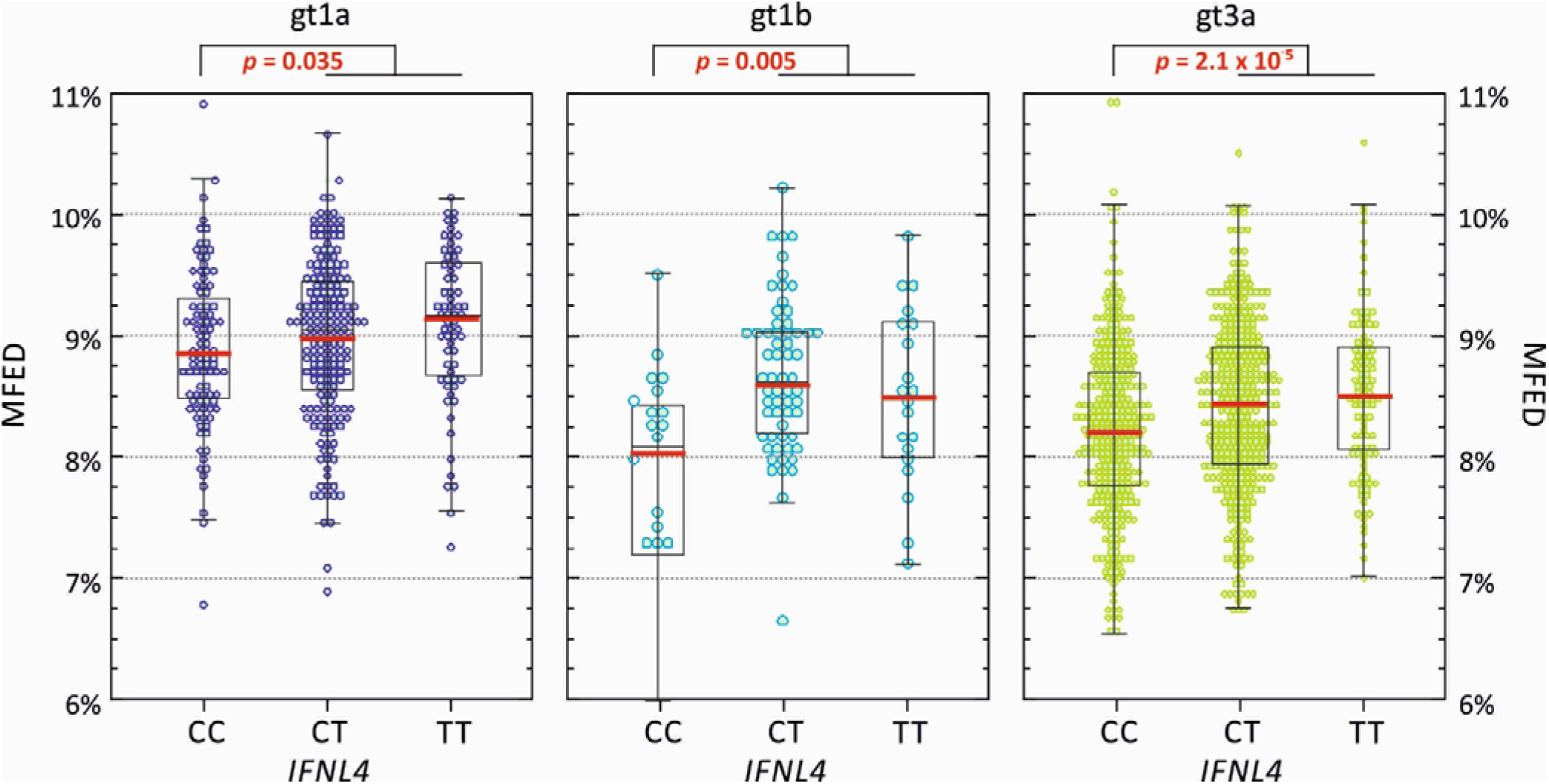
MFED VALUES OF gt1a, gt1b AND gt3a GENOMES RECOVERED FROM STUDY SUBJECTS WITH DIFFERENT *IFNL4* GENOTYPES. MFED Values for gt1a, gt1b and gt3a full genome sequences divided into the *IFNL4* genetic background of the host. The box plots show (from the top): 2 SDs above mean, 1 SD above mean, mean (red bar), 1 SD below mean and 2 SDs below mean. Differences in the distribution of MFED vales were calculated using the Kruskall-Wallace non-parametric test.

**FIGURE 7.**
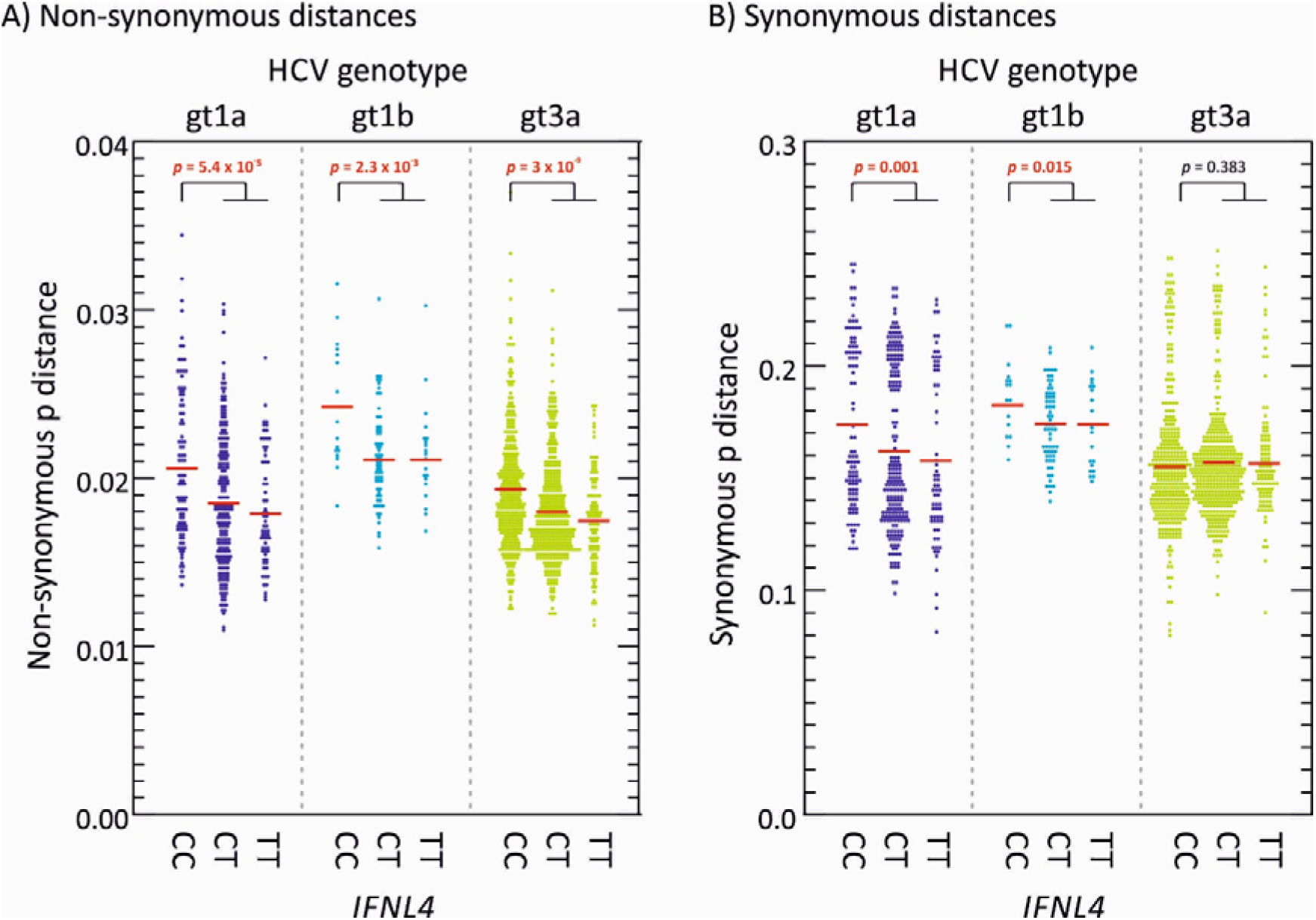
NON-SYNONYMOUS AND SYNONYMOUS SITE SEQUENCE DIVERGENCE OF POLYPROTEIN SEQUENCES OF DIFFERENT HCV GENOTYPES. Sequence distances between each study polyprotein sequence of gts 1a, 1b and 3a from a reconstructed consensus (ancestral) sequence of each genotype at (A) non-synonymous, and (B) synonymous sites. Red bars show mean values; differences in the distribution of MFED values were calculated using the Kruskall-Wallace test; significant values shown in red bold type.

**FIGURE 8.**
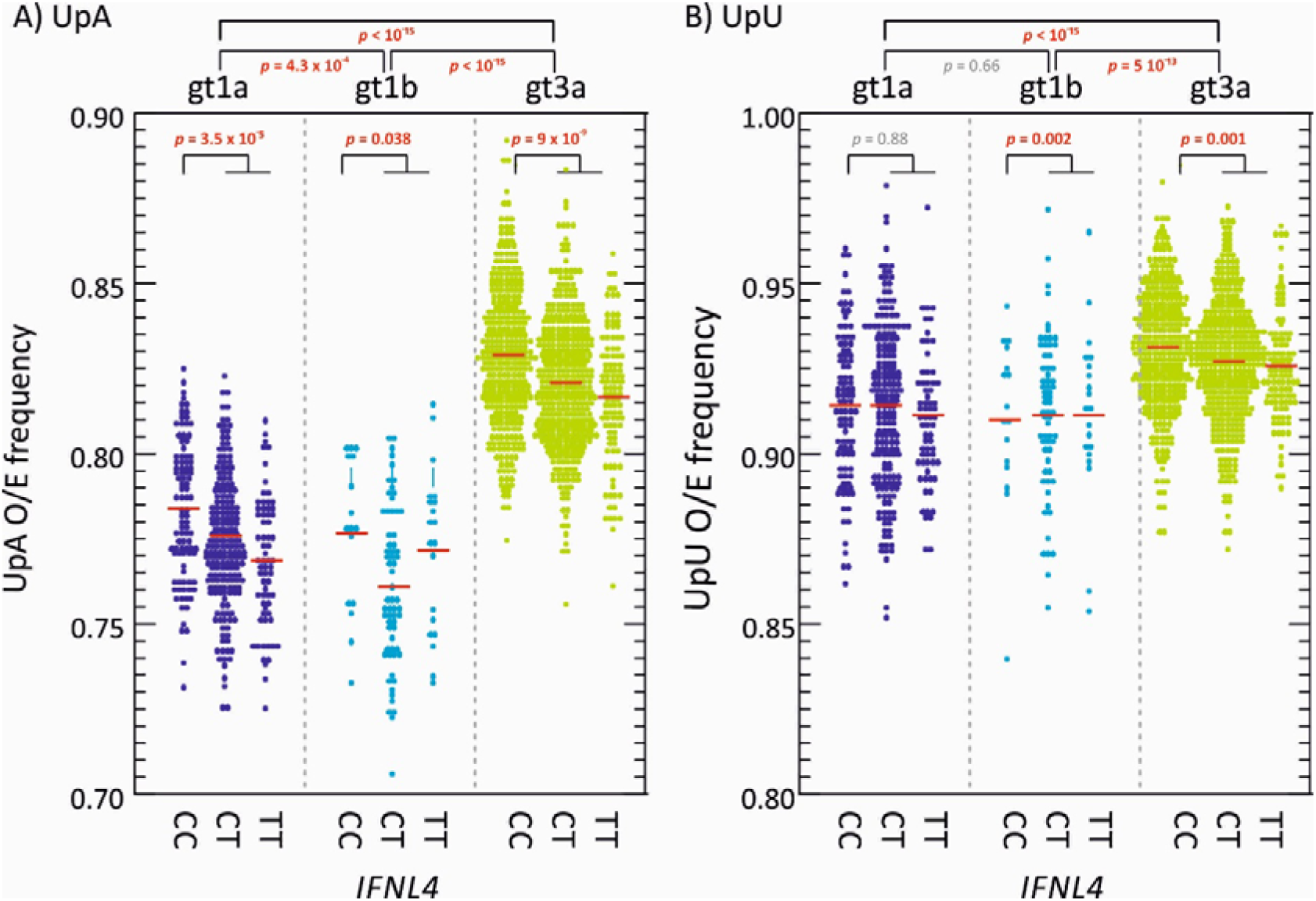
DISTRIBUTIONS OF UpA AND UpU FREQUENCIES. Observed to expected frequencies of (A) UpA, and (B) UpU dinucleotides targeted by RNAseL calculated for polyprotein sequence of gts 1a, 1b and 3a and different IFNL4 alleles. Red bars show mean values; differences in the distribution of O/E ratios were calculated using the Kruskall-Wallace test; significant values shown in red bold type.

The previously reported difference in UpA dinucleotide frequencies between *IFNL4* SNP genotypes in HCV genotype 3a (Ansari *et al.*, 2019) was reproduced in the larger datasets used in the current study for all 3 genotypes, although there was little systematic difference in UpU frequencies between different alleles (Fig. 8). No differences in CpG frequencies were identified (data not shown).

To determine whether population geographic differences of HCV influenced the association between *IFNL4* SNP rs12979860 with MFED values and UpA frequencies, we reanalysed associations in genotype 3a using the BOSON cohort, which includes study subjects from the UK, USA, Canada, Australia and New Zealand. As controls, we selected 500 SNPs from across the human genome that were frequency matched to *IFNL4* SNP rs12979860 (listed in Table S2; suppl. Data – avoiding X and Y chromosomes and SNPs within 200 kb from the *IFNL4* gene on chromosome 19). Linear regression was to test for associations between MFED and UpA frequency for each SNP and t-statistics for effects plotted for all 500 SNPs and rs12979860 (Fig. 9). We observed a normal distribution of t-statistics for the control SNPs, consistent with an absence of associations with MFED and UpA frequencies while accounting for possible effects of population structure. Contrastingly, the t-statistic for *IFNL4* SNP rs12979860 with MFED and UpA frequencies lies far outside the null distribution.

**FIGURE 9.**
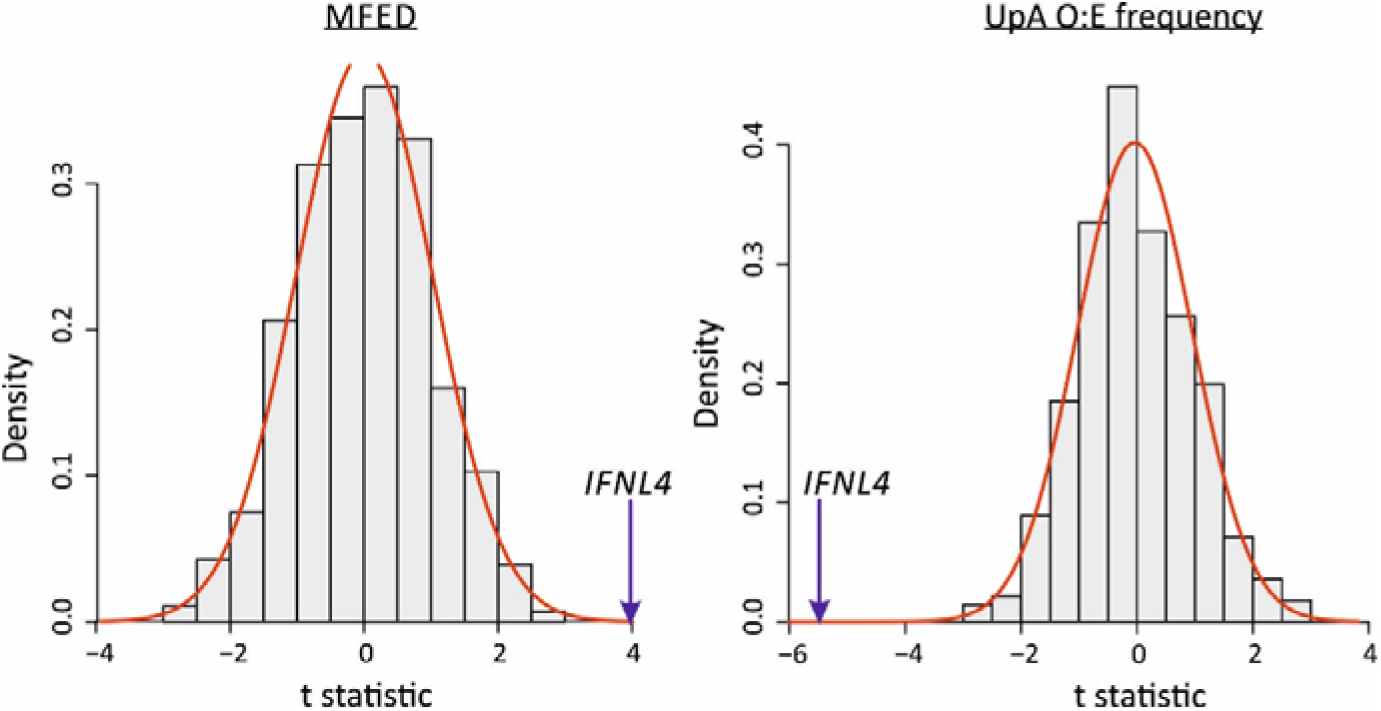
Association of MFED values and UpA frequencies with the *IFNL4* SNP rs12979860. The distribution of the t-statistics for 500 SNPs from across the genome frequency matched to the *IFNL4* SNP rs12979860 (SNPs listed in Table S2; Suppl. Data). The red line shows the fitted normal distribution to the data. The t-statistic is from a linear regression where the dependant variable is the MFED or UpA frequency and the independent variable is the SNP with a dominant genetic model (coded the same as *IFNL4* SNP rs12979860 CC vs. non-CC genotypes). The blue arrows indicate the t-statistic for the *IFNL4* SNP rs12979860 CC vs. non-CC genotypes.

## DISCUSSION

### Visualisation methods for RNA secondary structure

A diagrammatic representation of an individual stem-loop and its potential kissing loop / pseudoknot tertiary pairings represent the standard means for depicting RNA secondary and tertiary structures. Their utility, however, is limited as a means to incorporate information on RNA structural variability, such as different identities of the bases paired and their relative positions within a structure. RNA structure diagrams become increasingly difficult to construct for longer sequences, such as whole virus genomes analysed in the current study. To address these restrictions in our study of RNA structural diversity, we used a form of 3-dimensional representation of multiple sequences in alignments spanning the lengths of viral genomes. The depiction of stem-loop tips at a standard height (0) highlights the principal structural elements of an RNA folding prediction for each sequence while the size of the stem-loop is depicted of depth. In the current study, we colour-coded the StructureDist output but the 3-dimensional coordinate file produced by the program can be used for alternative forms of representation, such as 3-dimensional surface visualisations combined with object rotations.

The advantages of this form of whole genome representation is apparent when comparing predicted RNA structures for HCV genotypes 1-3, other viruses with GORS and viruses with unstructured genomes (Fig. 2). For the latter, the plethora of arbitrary, short and variable pairings in alignments of unstructured RNA sequences appear entirely distinct from the ordered sets of stem-loops visualised for those that possess sequence order-dependent pairings (high MFED values). The ability to display both pairing and conservation at each position in the genome provides a highly effective way to visualise which elements and structures are conserved across a dataset and the extent to which RNA structures may systematically vary between genotypes or serotypes. The calculation of a heterogeneity score for pairing predictions between individual sequences clearly distinguishes between viruses with unstructured and structured genomes (Fig. 5).

The accuracy of the depictions of RNA structure in contour plots is obviously only as good as the underlying algorithm used to predict RNA structure pairings. The analysis described in the current study used RNAFold, a widely used and validated structure prediction method based upon folding energy minimisation (Lorenz *et al.*, 2011). The underlying algorithm outperforms other energy minimisation algorithms in terms of speed and accuracy for structure prediction (Lorenz *et al.*, 2011) and is able to generate numeric characteristics of RNA folds, such as MFE values that are used in MFED calculations (Fig. 1). For contour plots, the accuracy of the predictions was increased through generating an ensemble of sub-optimal folds for each sequence fragment (Wuchty *et al.*, 1999) and deriving a consensus prediction based on a simple majority rule (>50%). Sites with multiple conflicting predictions, typically those in regions without organised RNA folding, were therefore excluded from the plots and sharpen the differentiation between structured and unstructured regions (Fig. 4). The calling framework in SSE can however be adapted to future bioinformatic developments in RNA structure prediction, including the use of algorithms that address the knotty problem of tertiary structure prediction (eg. (El Fatmi *et al.*, 2019; Jabbari *et al.*, 2018; Singh *et al.*, 2019)). It is entirely possible, for example, that pseuoknot or kissing loop interactions may play further stabilising roles in GORS-associated RNA structures. The perplexing spread of MFED values based only on secondary structure-based energy calculations within individual HCV subtypes (Fig. 1B) or genotypes (Fig. S4; Suppl. Data) might conceivably be reconciled if additional tertiary elements involved in RNA structure formation could be incorporated in MFE calculations.

### RNA structure plasticity

This exploratory study investigated the extent and conservation of RNA structure formation in different genotypes of HCV. In areas of the coding sequences where conserved pairing were predicted between genotypes, such as in the core and NS5B regions, these were fully consistent with those previously determined by nuclease mapping, SHAPE analysis and functional studies (Diviney *et al.*, 2008; Mauger *et al.*, 2015; McMullan *et al.*, 2007; Pirakitikulr *et al.*, 2016; Tuplin *et al.*, 2004; You *et al.*, 2004). The contour plots, however, revealed far more RNA structure that differed between genotypes throughout the E1-NS5B regions. Most RNA structures were indeed unique to individual genotypes of HCV (Fig. 2A) that otherwise shared only in their elevated MFED scores as a marker of RNA structure formation (Fig. 1B). These findings are consistent with the evident differences in RNA structures revealed by SHAPE mapping of genomes of genotypes 1a, 2a and 3a (Mauger *et al.*, 2015), and indeed visually apparent from the match between contour plot representations of the published SHAPE structures and RNAFold predictions for these genotypes (Fig. 3A). HCV therefore demonstrates considerable plasticity in the nature of its RNA pairings between subtypes and genotypes in most of the genome. This suggests that it is simply folding rather than the actual topography and potential interactions with cellular or viral RNA structures or proteins that is functionally important.

The highly variable structures formed by different genotypes visualised in the contour plots (Fig. 2A) account for the previous difficulties in mapping and functionally characterising RNA structures in HCV. Indeed, few of the RNA structures investigated functionally after their identification by SHAPE were conserved across genotypes and even fewer possessed obvious replication functions. For example, effects on virus replication kinetics were modest or absent in mutants with disrupted base pairings in SL783 (core gene), SL1412 (E1), SL6038 (NS4A) and SL8001 (NS5B) in the JFH genotype 2a strain (Pirakitikulr *et al.*, 2016). Similarly, disruption of J750 (core) and J8640 (NS5B) showed no effect on the replication of Jc1, while less than one log reductions were observed on disruption of J7880 (NS5B) and J8880 (NS5B) (Mauger *et al.*, 2015). Systematic, large scale mutagenesis of 17 consecutive sequence segments showed mild to modest replication effects only in the two final segments spanning positions 8441-8767 and 8768-9087 upstream of the CRE (SL9266) and an absence of cell culture functional elements in the rest of the coding region (Chu *et al.*, 2013). These effects are collectively different from the previously documented lethal phenotype associated with disruption the HCV CRE (SL9266 or NS5B 3.2) on replication that is conserved across all genotypes.

The diversity of predicted RNA structure elements between HCV variants (Fig. 2A; Fig. S3; Suppl. Data) was mirrored by FMDV. Its genome similarly shows elevated MFED scores across its genome (Fig. S4; Suppl. Data) despite the evident differences in RNA structures formed by different serotypes across its more variable structural gene region. Collectively, these findings challenge the prevailing paradigm of viral RNA structures being discrete elements, conserved in base-pairings with defined functions and being highly evolutionarily stable. GORS, based on the current and previous analyses, differs in all of these characteristics, being pervasive throughout the genome, variable in pairings, likely mediating a general conformational effect on interactions with the cell and being highly evolutionarily plastic. While it has been argued that many or most of RNA structures predicted in the coding region of Jc1 (genotype 2a) strain may play replication or regulatory roles (Pirakitikulr *et al.*, 2016), it seems difficult to imagine how any form of equivalence in functional properties could be maintained in other genotypes with quite radically different RNA structural organisations.

### The evolution of RNA structure

The timescale for the evolution of different HCV genotypes and subtypes is currently uncertain. However, its estimated nucleotide substitution rate of 5 – 10 × 10^−4^ substitutions per site per year (Gray *et al.*, 2011; Markov *et al.*, 2009) indicates that diversification of subtypes within a genotype has occurred over many hundreds of years; for example, the various genotype 2 subtypes have been proposed to have diverged in Guinea-Bissau in 1470 (range 1414-1582) (Markov *et al.*, 2009). The eight HCV genotypes may have originated proportionately earlier (Smith *et al.*, 1997). Given the evident differences between contour plots for HCV 1a, 1b, 2a and 3a (Fig. 3), HCV has evidently largely re-modelled its RNA secondary structure throughout the coding region over this period. At the same time, HCV has presumably managed to preserve sequence order-dependent RNA structure in its evolutionary intermediates, given the universal presence of elevated MFED values in all HCV genotypes (Fig. 1A; Fig. S4, Suppl. Data). On a more immediate evolutionary timescale, we have obtained evidence for the evolution of RNA structure over the course of infection. The emergence of systematic differences in MFED values between study subjects with different *IFNL4* genotypes (Fig. 7) indicates effects of interferon lambda expression on genome configurations of genotypes 1 and 3 that could only have arisen over the perhaps 20-40 years course of infection within each individual.

A link between GORS and cellular responses to infection was previously inferred from the evident association between its presence and virus persistence (Davis *et al.*, 2008; Simmonds *et al.*, 2004). It has been proposed that RNA structure formation may assist in the evasion of immune recognition of genomic RNA by RNAseL and PKR during persistent infections (Mauger *et al.*, 2015). RNAseL is activated by 2⍰–5⍰ oligoadenylate molecules (2-5A) produced by a range of oligoadenylate synthetases (OASs) and subsequently targets single stranded RNA sequences for cleavage at UpA and UpU dinucleotide sites (Han and Barton, 2002; Wreschner *et al.*, 1981). The mammalian OAS / RNAseL plays a major role in the control of HCV and other RNA virus infections (Chakrabarti *et al.*, 2011; Han and Barton, 2002; Kristiansen *et al.*, 2011; Li *et al.*, 2016; Silverman, 2007) and may significantly influence the outcomes of HCV infections and their responsiveness to treatment. Differences between RNAseL sensitivity between HCV genotypes and associated differences in UpA frequencies (Washenberger *et al.*, 2007)(Fig. 8) and RNA structure formation (Fig. 1B) are not inconsistent with this association.

In unravelling these factors, it has been found by several groups that intrinsic levels of ISG expression are greater in genotype 1 infections than in genotype 2 or 3 (Chen *et al.*, 2010; d’Avigdor *et al.*, 2019; Holmes *et al.*, 2015). The internal environment in genotype 1-infected hepatocytes may be consequently more hostile for HCV replication. Among the various antiviral pathways activated by IFN, RNAseL potently restricts RNA virus replication through cleavage of viral RNA genomic sequences at UpA and UpU dinucleotide sites. Significantly, HCV genotype 1 shows a markedly greater suppression of UpA frequencies than genotypes 2 and 3 and consequent alterations in codon usage (Han and Barton, 2002; Washenberger *et al.*, 2007) (also evident in Fig. 8); this has been proposed by the authors as a necessary adaptive change to minimise the antiviral activity of RNAseL. The greater degree of folding of RNA in genotype 1 (mean MFED values of 9.0%, compared to 8.4% in genotype 3 (Fig. 1B) may similarly reflect an adaptive change to reduce the frequencies of UpA/UpU dinucleotides in single stranded RNA contexts that can be targeted by RNAseL.

The same pattern of linked variables is mirrored on a smaller scale between individuals with different *IFNL4* genotypes. The greater degree of RNA structure formation in those with CT and TT genotypes (Fig. 1B) and the reduced frequencies of UpA dinucleotides (Fig. 8)(Ansari *et al.*, 2019) represents a microcosm of the larger virus genotype effects. They may reflect the much shorter term outcomes of more effective virus control mediated through RNAseL and other ISGs consequent to IFNλ4 expression in the liver during the course of chronic infection (Dill *et al.*, 2011). Extending the model further, the reported evidence for less divergence of HCV genotype 3 sequences in those with CT or TT genotypes (Ansari *et al.*, 2019) which was also reflected in genotypes 1a and 1b (Fig. 7) may reflect the greater selection strength operating on viruses in cells with enhanced innate immune control. The finding in the current study that diversity at synonymous sites is also more restricted in non-CC individuals indicates a selection pressure potentially operating at the RNA level rather than on the viral proteome as previously suggested (Ansari *et al.*, 2019). The observation in the latter study for associations at synonymous positions within several codons with *IFNL4* SNP rs12979860 provides further evidence for selection at the RNA level. Its focus on genotype 3a, where the differences in synonymous variability between CC and non-CC individuals is much less apparent than in genotypes 1a and 1b (Fig. 7) potentially accounts for the differences in conclusions reached. RNA structure requirements and maintaining low frequencies of UpA dinucleotides may indeed place quite different selection pressures of HCV infection in different *IFNL4* backgrounds at both non-synonymous and synonymous sites. More broadly, the cellular host response may therefore be a potent factor influencing the course of evolution of HCV within an infected individual.

To conclude, we have shown that RNA secondary structure in HCV genomes pervades whole genomes of HCV and other viruses that establish persistent infections. GORS not only shows substantial changes in response to the host environment mediated through the *IFNL4* polymorphism, but it has also been entirely remodelled in most of the genome over the longer period of its divergence into different subtypes and genotypes (Fig. 2). In contrast to discrete, conserved RNA structural elements used by RNA viruses for replication and translation functions, GORS is pervasive and profoundly adaptive over even relatively short evolutionary periods. While the current study findings support the previously suggested link between RNA folding and RNAseL susceptibility, the broader underlying reasons for virus genomes becoming structured in this way require considerable further investigation.

## MATERIALS AND METHODS

### Sequence datasets

HCV complete polyprotein sequences were obtained from previously recruited cohorts (BOSON, the Early Access Programme [EAP], STOP-HCV-1 and the UK cirrhosis study) (Fawsitt *et al.*, 2019; Foster *et al.*; McLauchlan *et al.*, 2017). Patients from the BOSON cohort originated from Australia, Canada, New Zealand, United Kingdom and United States, while the remaining cohorts were of UK-only origin. The BOSON study was conducted in accordance with the International Conference on Harmonisation Good Clinical Practice Guidelines and the Declaration of Helsinki (clinical trial registration number: NCT01962441). The EAP study conformed to the ethical guidelines of the 1975 Declaration of Helsinki as reflected in a priori approval by the institution’s human research committee. The EAP patients were enrolled by consent into the HCV Research UK registry. Ethics approval for HCV Research UK was given by NRES Committee East Midlands - Derby 1 (Research Ethics Committee reference 11/EM/0314). A subset of genotype 3a subjects with self-reported white ancestry infected with HCV genotype 3a for which we had obtained both host genome-wide SNP data was selected for genome wide association study. Duration of infections were not determined for the study subjects although the cohorts varied in their progression and disease severity, ranging from mild (STOP-HCV-1, BOSON) to severe (EAP, cirrhosis cohort).

Available coding complete genome sequences of other viruses (FMDV serotype 0 [FMDV-O], HPgV-1, HPeV-3, JEV, MNV and EV-A71 were downloaded from Genbank in 2019; redundant sequences were removed by filtering out sequences showing <1% nucleotide sequence from others in the dataset (accession numbers of the selected sequences listed in Table S1; Suppl. Data).

### RNA structure prediction

Minimum folding energies were predicted using the RNAFold.exe program in the RNAFold package, version 2.4.2 (Lorenz *et al.*, 2011) using with default parameters (37C, allow isolated base pairs, linear sequence, unlimited pairing span) using sequential 240 base sequence fragments incrementing by 12 bases between fragments. Contour plots were generated from ensemble RNA structure predictions (Wuchty *et al.*, 1999) of sequential 1600 base fragments of the alignment incrementing by 400 bases between fragments using the program SubOpt.exe, with the following additional parameters - all sub-optimal structures, maximum 50 analysed. Pairing predictions supported by >50% of sub-optimal structures were used in the consensus contour plot; predictions from sites without a majority prediction were discarded. Folding heterogeneity was expressed as the difference in folding depth between different sequences at each position in the alignment; low values therefore correspond to sites with concordant pairing predictions. Analyses using RNAFold.exe and SubOpt.exe were invoked through the programs Folding Energy Scan and StructureDist in the SSE package version 1.4 (Simmonds, 2012) (http://www.virus-evolution.org/Downloads/Software/). StructureDist in version 1.4 of SSE has been extended the calculation of folding depths used for contour plotting. Sequence distances were calculated in the SSE package.

### Host genetics

Host genetics information was based on host genome-wide genotyping platform (Affymetrix UK Biobank Chip) and other demographic data collected for previous studies (Ansari *et al.*, 2019; Ansari *et al.*, 2017).

## ACKNOWLEDGEMENTS

STOP-HCV Collaborators: Ball J, Brainard D, Burgess G, Cooke GS, Dillon J, Foster G, Gore C, Guha N, Halford R, Whitby K, Holmes C, Howe A, Hudson E, Hutchinson S, Khakoo S, Klenerman P, Martin N, Massetto B, Mbisa T, McHutchison J, McKeating J, Miners A, Murray A, Shaw P, Spencer C, Thomson E, Vickerman P, Zitzmann N.

