## Supplementary Data for "Impact of virus subtype and host *IFNL4* genotype on large-scale RNA structure formation in the genome of hepatitis C virus"

TABLE S1: GENBANK ACCESSION NUMBERS OF VIRUS SEQUENCES

FMDV-O: AY593824, JN998086, FJ461345, DQ248888, AY686687, EF552691, JX066664, JX040492, KF694737, AY593813, AY593822, KJ206910, KJ825804, AY593826, KF112882, HM191257, GU384683, HM008917, EU448369, AY593817, GU125647, AY593833, KF112880, KF112885, JX869177, HQ632769, AY593827, AF511039, AY333431, DQ478937, KF112888, FJ461344, AY593812, HQ412603, HQ113232, KJ206908, HQ632768, AY359854, FJ175664, KF112879, JQ900581, AY593830, AY317098, AY312588S2, GU125650, KF501486, HQ009509, AY593834, AF377945, DQ119643, KF112886, EU140964, HQ632770, KF501488, KM268895, HQ632771, JX869178, AY593825, JF749851, HQ632772, HQ268524, AY593823, AY593821, KF985189, KJ206909, JX040501, AY593828, AY593829, KJ825801, AB079061, AY593814,

HPgV-1 (n=100): AF121950, D87255, AF081782, D90600, LT009483, LT009487, LT009485, LT009494, AY196904, MH053119, LT009481, LT009478, HGU44402, LT009479, AF309966, MK291245, MK291244, AF031827, MH053115, KP259281, MH053120, HGU45966, MH053118, MH053121, HGU63715, LT009486, AB003289, LT009489, AF104403, JN127373, D90601, AB013501, MH053116, D87711, HGU36380, AB008335, D87263, D87712, D87713, KM670109, D87262, D87709, KM670096, LT009490, AB008342, KM670099, D87714, LT009480, KP710601, KM670110, LT009482, D87710, AB021287, KM670097, LT009484, KP710599, HQ331235, KM670100, HQ331234, LT009488, KM670107, HQ331233, KM670108, KP710604, KM670102, KM670106, KC618400, D87708, AB008336, KM670098, HGU94695, KP710605, KP710600, KC618401, MH746815, KC618399, AB013500, KP710598, KC618398, KM670101, MH053117, KP710602, KP710603, AB003288, KP710606, AB018667, HGU75356, AY949771, D87715, AB003290, AB003293, AF006500, AB003291, AB003292, MH179063, LT009493, MK291243, LT009492, LT009491, AF121950,

MNV (n=65): FJ446719, JN975495, EU004682, DQ223042, EU004668, AB435515, EU004680, JF320653, JN975498, EU004675, JF320647, DQ223043, EU004671, FJ446720, EU004674, EU004670, EU004681, EU004658, EU004677, HQ317203, EU004667, JN975493, JF320652, EU854589, JN975496, EF531290, AB601769, JN975494, JQ237823, EU004678, EU004663, DQ911368, EF531291, EU004683, EU004679, EU004664, DQ223041, EU004673, DQ285629,

EU004656, EU004659, EU004661, EU004662, AY228235, EU004655, EF014462, EU004654, EU004657, JF320648, JF320649, JF320650, JF320645, JF320651, JF320646, EU004669, EU004665, EU004672, EU004660, AB435514, JF320644, EU004676, JQ658375, EU004666,

FMDV-O (n=67): AY593824, JN998086, FJ461345, DQ248888, AY686687, EF552691, JX066664, JX040492, KF694737, AY593813, AY593822, KJ206910, KJ825804, AY593826, KF112882, HM191257, GU384683, HM008917, EU448369, AY593817, GU125647, AY593833, KF112880, KF112885, JX869177, HQ632769, AY593827, AF511039, AY333431, DQ478937, KF112888, FJ461344, AY593812, HQ412603, HQ113232, KJ206908, HQ632768, AY359854, FJ175664, KF112879, JQ900581, AY593830, AY317098, AY312588S2, GU125650, KF501486, HQ009509, AY593834, AF377945, DQ119643, KF112886, EU140964, HQ632770, KF501488, KM268895, HQ632771, JX869178, AY593825, JF749851, HQ632772, HQ268524, AY593823, AY593821, KF985189, KJ206909, JX040501, AY593828, AY593829, KJ825801, AB079061, AY593814,

HPeV-3(n=38): KX068679, KM986843, KJ659490, AB084913, AJ889918, KY020128, JX682576, GQ183029, GQ183033, GQ183032, GQ183031, GQ183026, GQ183027, KY556675, MK604044, MF371334, MK604049, KT879915, KT879922, KY351616, KY351618, JX826607, MF797924, MF797928, MF797929, KT626009, MF797935, MF797926, MF797925, MF797932, MF797916, MF797931, AB668029, LC129275, LC129276, LC129274, AB759205, AB759199,

JEV (n=65): JN381838, KT957421, JN381872, JN381830, JQ086763, JN381834, AF075723, HQ652538, AB241119, JN381867, AY184212, AB551990, AB569988, AF217620, KM658163, AB471669, AY303796, AB051292, KY927815, KF297916, HQ223286, M18370, JF706283, JN381853, AF080251, KY927818, AY316157, GQ902063, JF915894, KY078829, JN381863, AB551991, AF045551, KT229572, L48961, HQ223287, JN381843, JF706270, HQ223285, GQ902062, GQ902061, KM677246, KT957420, AY303794, KT229575, JF706279, HE861351, JN381868, JN381870, KY927816, U47032, AF098736, EF623987, JN381873, AB241118, JN381848, JN381866, AF221500, AY303792, GU205163, KF667311, GQ902060, JN381851, KU351667, EU429297,

EV-A71 (n=238): MK652139, DQ341367, MG976581, KX372324, JX244184, MH716347, JQ639384, LT719065, LC506514, KC436271, MG214681, KT428646, KP289432, MH716380, JF738002, HQ647173, GQ279369, KU936120, HM807310, FJ607337, FJ357384, MG367595, LC321989, MH716337, AB575917, KX372322, KJ004560, KF982854,

JQ074189, MG367594, EU131776, LT719063, LT719067, AB575918, MN254979, MF662695, KJ400360, U22522, KJ686246, KU936132, AB575938, HM002487, HQ647172, KJ686249, JN992284, KC436265, AB575935, KU936128, FJ194964, KJ784495, AB747375, JX678885, DQ341358, LC375764, AB575928, MF973167, LC506513, FJ357376, LK985324, MG672479, KF142413, MG773126, KX372317, KP289419, KP289425, KP289422, AB575911, KT345959, MG672481, GQ994992, DQ341364, DQ341357, KU936121, KC436266, KY612315, KC109780, AB575914, GQ231939, MH111073, JQ681218, KJ686308, MH716310, AF302996, GQ994989, KY315729, AM396584, LT719064, KM055005, KU254596, KP308450, HQ456306, KC436270, MG367600, KU936125, JQ742001, KC954664, FJ607336, HQ189392, KT008669, DQ341355, JQ742002, KX372328, JQ280307, KX372310, KM673244, DQ341359, KT428647, AM396587, KJ686262, FJ606447, KP289431, KX197462, AB575936, MG672478, LT719068, KP289424, EF063152, MG207962, EU414333, FJ360545, AF316321, JF738001, MF662701, KJ686208, KP308426, JF799986, MH484070, KF974790, GQ231934, U22521, AB575923, GQ994991, MG431943, JQ316638, DQ341360, KX372308, AY465356, HQ647179,

HCV all genotypes (n=125): EF108306, AF009606, D90208, D14853, KJ439768, KC248194, AM910652, KC248198, KJ439772, KC248193, KJ439778, KJ439781, KJ439774, HQ537007, AJ851228, KC248195, KJ439780, KJ439779, KJ439776, KJ439777, D00944, D10988_D01221, D50409, JF735114, JF735120, KC844042, DQ155561, HM777358, AB031663, JF735111, FN666428, JF735115, KC197238, JF735112, JF735116, JF735118, JF735117, JF735119, JF735110, KC197236, KC197237, KC197239, D17763, D49374_D26556, KJ470619, KJ470618, JX227954, JF735121, FJ407092, D63821, JF735124, Y11604, FJ462436, DQ418786, EF589161, FJ462432, EU392173, FJ839870, FJ462433, FJ462441, FJ462440, FJ462431, FJ462434, FJ462439, JF735136, FJ839869, HQ537009, HQ537008, FJ025855, FJ025854, JX227964, JF735127, JF735132, JF735131, JF735130, JF735129, JF735138, JF735135, JF735134, AF064490, Y12083, D84262, EF424629, D84263, DQ314805, DQ835760, D63822, D84265, DQ835770, DQ835761, D84264, EF424628, DQ835767, DQ278894, EF424627, EF424626, EF424625, EU408328, EU408329, EF632071, EU246940, EU158186, DQ278892_AY834954_AY835335, EU643836, EU408330, JX183552, KJ567651, KM252789, JX183557, DQ278891, DQ278893, JX183558, JX183553, JX183554, JX183551, JX183549, JX183550, KJ470620, KJ470621, KJ470622, KJ470623, KJ470624, KJ470625, KC844039, KC844040,

TABLE S2

LIST OF CONTROL SNPs

| Chr | Position | rs_ID | Orient |
|---|---|---|---|
| chr4 | 12983343 | rs10000091 | + |
| chr1 | 111734975 | rs1000947 | + |
| chr4 | 63197594 | rs10010596 | + |
| chr4 | 100695124 | rs10030146 | + |
| chr4 | 183534161 | rs10033074 | + |
| chr4 | 114323981 | rs10034750 | + |
| chr9 | 34631162 | rs10046843 | + |
| chr4 | 23288098 | rs10050029 | + |
| chr8 | 26285479 | rs10093390 | + |
| chr8 | 28228291 | rs10094977 | + |
| chr20 | 38732301 | rs1010321 | + |
| chr8 | 60948037 | rs10108543 | + |
| chr8 | 118714003 | rs10113233 | + |
| chr9 | 108381380 | rs10115131 | + |
| chr7 | 143452257 | rs1014069 | - |
| chr14 | 44561671 | rs10149946 | + |
| chr2 | 32479118 | rs10182170 | + |
| chr2 | 151625463 | rs10186482 | + |
| chr2 | 235032893 | rs10190447 | + |
| chr7 | 34445525 | rs10241710 | + |
| chr7 | 151127257 | rs10256812 | + |
| chr7 | 50273756 | rs10276619 | + |
| chr6 | 24961163 | rs1028477 | - |
| chr10 | 130892388 | rs1036270 | + |
| chr22 | 37283722 | rs1041895 | + |
| chr6 | 55145701 | rs10456701 | + |
| chr5 | 167421729 | rs10475843 | + |
| chr5 | 147809331 | rs10477350 | + |
| chr7 | 142469229 | rs10487531 | + |
| chr9 | 27468463 | rs10511816 | - |
| chr9 | 87555970 | rs10512187 | + |
| chr12 | 125025558 | rs1055472 | + |
| chr1 | 84647546 | rs1057746 | - |
| chr10 | 78299615 | rs10762779 | + |
| chr12 | 125726715 | rs10773198 | + |
| chr11 | 134901914 | rs10791403 | + |
| chr10 | 7904008 | rs10795572 | + |
| chr1 | 30105771 | rs10799077 | + |
| chr7 | 129977887 | rs10808241 | + |
| chr9 | 34596680 | rs10814125 | + |
| chr11 | 48560213 | rs10838956 | + |
| chr12 | 3218420 | rs10848829 | + |
| chr12 | 80057464 | rs10862038 | + |
| chr1 | 9388590 | rs10864413 | + |
| chr9 | 69204747 | rs10870017 | + |
| chr10 | 112662640 | rs10885369 | + |
| chr10 | 117770266 | rs10886131 | + |
| chr1 | 55292134 | rs10888912 | + |
| chr11 | 115233890 | rs10891814 | + |
| chr5 | 139396890 | rs10900862 | + |
| chr12 | 131620248 | rs10902530 | + |
| chr10 | 123066903 | rs10902870 | + |
| chr10 | 14671222 | rs10906740 | + |
| chr10 | 18964758 | rs11007866 | + |
| chr10 | 128962749 | rs11016572 | + |
| chr11 | 48761945 | rs11040016 | + |
| chr17 | 6374568 | rs11078621 | + |
| chr21 | 25965946 | rs11087985 | + |
| chr12 | 95596979 | rs11108147 | + |
| chr1 | 210824120 | rs11119611 | + |
| chr20 | 23343350 | rs1112819 | + |
| chr16 | 74943972 | rs11149785 | + |
| chr1 | 40515059 | rs11208299 | + |
| chr2 | 18902959 | rs1122624 | - |
| chr8 | 10393755 | rs11249999 | + |
| chr10 | 6288034 | rs11257854 | + |
| chr12 | 122327956 | rs1129167 | - |
| chr11 | 134294724 | rs1144221 | - |
| chr14 | 55845400 | rs11621920 | + |
| chr14 | 104494153 | rs11626836 | + |
| chr15 | 73883936 | rs11639300 | + |
| chr16 | 88078091 | rs11639845 | + |
| chr16 | 31410096 | rs11640148 | + |
| chr16 | 51524634 | rs11646208 | + |
| chr17 | 4453080 | rs11655342 | + |
| chr2 | 18338468 | rs11689614 | + |
| chr4 | 168047067 | rs11728448 | + |
| chr4 | 16994880 | rs11731316 | + |
| chr4 | 179752670 | rs11736914 | + |
| chr4 | 187969824 | rs11737485 | + |
| chr5 | 50616675 | rs11746825 | + |
| chr6 | 70102381 | rs11759810 | + |
| chr7 | 12556990 | rs11764174 | + |
| chr1 | 157163842 | rs1176544 | + |
| chr8 | 122795139 | rs11775943 | + |
| chr10 | 117148539 | rs11819305 | + |
| chr14 | 103970594 | rs11851097 | + |
| chr16 | 80669860 | rs11863297 | + |
| chr9 | 84701953 | rs1187336 | - |
| chr3 | 193956972 | rs11914490 | + |
| chr3 | 172584145 | rs11922343 | + |
| chr1 | 220500441 | rs12120994 | + |
| chr1 | 34325265 | rs12130968 | + |
| chr1 | 91570783 | rs12141173 | + |
| chr21 | 21147102 | rs12152089 | + |
| chr2 | 212735163 | rs12185565 | + |
| chr6 | 44494493 | rs12192702 | + |
| chr1 | 247438293 | rs12239046 | + |
| chr12 | 16124240 | rs12297516 | + |
| chr14 | 49490614 | rs12323543 | + |
| chr11 | 4848419 | rs12361955 | + |
| chr11 | 48534405 | rs12365220 | + |
| chr14 | 53694533 | rs12433844 | + |
| chr17 | 37137586 | rs12450937 | + |
| chr3 | 169602805 | rs12487498 | + |
| chr6 | 140866118 | rs12524925 | + |
| chr6 | 55479176 | rs12526863 | + |
| chr7 | 118757867 | rs12536552 | + |
| chr8 | 80421136 | rs12545933 | + |
| chr9 | 80192588 | rs12554086 | + |
| chr15 | 67103580 | rs12594610 | + |
| chr1 | 214836031 | rs1261868 | - |
| chr4 | 61235270 | rs12639891 | + |
| chr5 | 175865617 | rs12652846 | + |
| chr6 | 144202524 | rs12664476 | + |
| chr8 | 135820613 | rs12678843 | + |
| chr7 | 113677653 | rs12705891 | + |
| chr12 | 14510823 | rs12708367 | + |
| chr1 | 190021748 | rs12732394 | + |
| chr1 | 38962356 | rs12733433 | + |
| chr3 | 59676390 | rs1281463 | + |
| chr12 | 94754980 | rs12822392 | + |
| chr15 | 54465639 | rs12904540 | + |
| chr17 | 72996427 | rs12947636 | + |
| chr18 | 45660009 | rs12957040 | + |
| chr2 | 119112657 | rs13009596 | + |
| chr21 | 45347113 | rs13049248 | + |
| chr3 | 194698992 | rs13084683 | + |
| chr4 | 101210521 | rs13130412 | + |
| chr4 | 142689140 | rs13133181 | + |
| chr5 | 154408936 | rs13162143 | + |
| chr5 | 12013356 | rs13171948 | + |
| chr5 | 29952825 | rs13183145 | + |
| chr5 | 3855257 | rs1319960 | + |
| chr8 | 71089696 | rs13251367 | + |
| chr8 | 126268783 | rs13259816 | + |
| chr8 | 18658330 | rs13274103 | + |
| chr9 | 32205706 | rs1331240 | + |
| chr16 | 74538674 | rs13336493 | + |
| chr2 | 170214402 | rs13385825 | + |
| chr13 | 87501880 | rs1340833 | + |
| chr22 | 49532624 | rs134452 | - |
| chr16 | 11585967 | rs1345441 | - |
| chr7 | 54349136 | rs1357884 | - |
| chr5 | 156956411 | rs1363232 | - |
| chr18 | 31207561 | rs1365294 | - |
| chr9 | 78928950 | rs1371013 | - |
| chr13 | 60391353 | rs1390157 | + |
| chr3 | 28917720 | rs1392307 | + |
| chr12 | 99476849 | rs1398084 | + |
| chr2 | 64558019 | rs1421337 | + |
| chr11 | 131030204 | rs1433982 | + |
| chr4 | 108843849 | rs1452688 | - |
| chr15 | 94981210 | rs1457851 | - |
| chr10 | 60314381 | rs1471246 | + |
| chr12 | 25065406 | rs1497249 | - |
| chr2 | 57476332 | rs1518685 | - |
| chr2 | 166335304 | rs1528487 | - |
| chr10 | 2397288 | rs1532215 | + |
| chr21 | 38987079 | rs1537106 | + |
| chr16 | 82529949 | rs1560824 | - |
| chr8 | 4844711 | rs1561360 | + |
| chr17 | 73242948 | rs1566290 | + |
| chr6 | 32701241 | rs1612904 | - |
| chr17 | 18930497 | rs1634411 | - |
| chr10 | 82125138 | rs1649934 | - |
| chr1 | 229171169 | rs16849500 | + |
| chr8 | 40639459 | rs16889787 | + |
| chr1 | 150319969 | rs1694379 | - |
| chr16 | 82642261 | rs16957935 | + |
| chr16 | 25067419 | rs16974327 | + |
| chr8 | 4247625 | rs1714746 | - |
| chr5 | 98319582 | rs17165849 | + |
| chr4 | 73595110 | rs17192483 | + |
| chr14 | 59251812 | rs17255451 | + |
| chr17 | 18937120 | rs1737944 | - |
| chr2 | 73695962 | rs17434655 | + |
| chr11 | 61888645 | rs174455 | + |
| chr15 | 77368003 | rs17470228 | + |
| chr10 | 29949908 | rs1756752 | - |
| chr15 | 51397575 | rs17648128 | + |
| chr15 | 51402295 | rs17648140 | + |
| chr14 | 38945690 | rs17670531 | + |
| chr21 | 41898747 | rs177933 | + |
| chr18 | 73625792 | rs17829298 | + |
| chr21 | 24122423 | rs1783355 | + |
| chr18 | 13116433 | rs1786263 | - |
| chr14 | 30858598 | rs179666 | - |

| | | | |
|---|---|---|---|
| chr7 | 44834514 | rs1802437 | - |
| chr2 | 236933481 | rs1821655 | + |
| chr3 | 103875711 | rs1824240 | + |
| chr5 | 58907240 | rs184187 | + |
| chr3 | 72500964 | rs1898634 | + |
| chr4 | 131514178 | rs1911889 | + |
| chr12 | 114268404 | rs1920591 | + |
| chr5 | 30870751 | rs1921111 | + |
| chr5 | 136700906 | rs193491 | - |
| chr10 | 109646537 | rs1936138 | - |
| chr11 | 98269834 | rs1941784 | - |
| chr7 | 29493778 | rs1946134 | + |
| chr2 | 227198074 | rs1950135 | - |
| chr14 | 32417836 | rs1957007 | + |
| chr6 | 54908016 | rs1963697 | - |
| chr6 | 7922058 | rs197113 | + |
| chr2 | 73700971 | rs2001490 | + |
| chr1 | 17071209 | rs2016693 | - |
| chr18 | 68729625 | rs2048329 | + |
| chr3 | 150914987 | rs2061201 | - |
| chr6 | 98946786 | rs2064146 | - |
| chr14 | 103412850 | rs2065018 | - |
| chr22 | 32435553 | rs2076050 | + |
| chr2 | 85668215 | rs2077079 | - |
| chr2 | 236945804 | rs2082870 | + |
| chr4 | 53564319 | rs2099453 | + |
| chr3 | 46561246 | rs2139633 | + |
| chr20 | 10847619 | rs2144186 | - |
| chr6 | 18207102 | rs214600 | - |
| chr10 | 118951825 | rs2148496 | - |
| chr1 | 241480284 | rs2153693 | + |
| chr7 | 32297831 | rs215606 | + |
| chr2 | 77924959 | rs2163875 | - |
| chr14 | 32491287 | rs2180866 | + |
| chr10 | 36774764 | rs2185863 | - |
| chr12 | 11702690 | rs2187642 | + |
| chr14 | 80016476 | rs2193131 | + |
| chr12 | 99234519 | rs2201160 | + |
| chr2 | 235335632 | rs2217538 | - |
| chr16 | 31363214 | rs2230429 | + |
| chr22 | 37066886 | rs2235321 | + |
| chr17 | 82675394 | rs2248371 | + |
| chr8 | 11853550 | rs2272767 | + |
| chr15 | 72735137 | rs2277598 | - |
| chr14 | 64796582 | rs229587 | + |
| chr1 | 206474694 | rs2297546 | - |
| chr1 | 119383313 | rs2298028 | + |
| chr11 | 20638211 | rs2298826 | + |
| chr16 | 661905 | rs2301426 | - |
| chr17 | 5532943 | rs2301582 | - |
| chr17 | 40935654 | rs2315602 | + |
| chr22 | 24602652 | rs2330805 | + |
| chr9 | 30077007 | rs2360147 | + |
| chr13 | 103960244 | rs2391059 | + |
| chr10 | 25750132 | rs2442855 | - |
| chr12 | 48793667 | rs2446990 | + |
| chr11 | 27379170 | rs2472625 | - |
| chr14 | 88267790 | rs2474952 | - |
| chr1 | 2205929 | rs2503712 | + |
| chr3 | 135050073 | rs253876 | - |
| chr5 | 172460777 | rs2569245 | + |
| chr5 | 75742893 | rs258494 | + |
| chr16 | 61089243 | rs2639889 | - |
| chr5 | 115842698 | rs26538 | + |
| chr5 | 115229045 | rs265436 | + |
| chr16 | 78547266 | rs2667649 | + |
| chr4 | 74011427 | rs2668076 | + |
| chr14 | 99180992 | rs2693680 | + |
| chr11 | 75147924 | rs2712806 | + |
| chr5 | 96384871 | rs271942 | - |
| chr5 | 60456813 | rs27223 | + |
| chr1 | 3122059 | rs2742672 | - |

| | | | |
|---|---|---|---|
| chr4 | 3718102 | rs2748777 | - |
| chr2 | 32557696 | rs2754513 | + |
| chr1 | 57218388 | rs2764660 | + |
| chr9 | 69463950 | rs2781538 | + |
| chr13 | 97682683 | rs2783482 | + |
| chr10 | 117436739 | rs2794424 | + |
| chr12 | 95428932 | rs280965 | + |
| chr1 | 72346757 | rs2815752 | - |
| chr5 | 75745736 | rs28225 | - |
| chr21 | 21133424 | rs2826674 | + |
| chr16 | 77555105 | rs282985 | + |
| chr5 | 142108414 | rs28372841 | + |
| chr12 | 132427238 | rs28386864 | + |
| chr6 | 62385346 | rs2842783 | - |
| chr6 | 31365526 | rs2844580 | - |
| chr12 | 132427669 | rs28451398 | + |
| chr20 | 57491781 | rs2865387 | + |
| chr5 | 51023481 | rs28654104 | + |
| chr7 | 212512 | rs28706381 | + |
| chr17 | 36887732 | rs2898660 | - |
| chr4 | 56179091 | rs2899056 | + |
| chr8 | 17155132 | rs2904659 | + |
| chr5 | 76272228 | rs2937715 | - |
| chr1 | 84582958 | rs2945152 | - |
| chr12 | 56720523 | rs2958127 | - |
| chr16 | 9153026 | rs2965895 | - |
| chr12 | 52424720 | rs298104 | - |
| chr6 | 56746538 | rs3002005 | + |
| chr3 | 11749236 | rs301554 | + |
| chr18 | 50448374 | rs3017261 | + |
| chr2 | 230982495 | rs3106082 | + |
| chr7 | 135702747 | rs3112346 | + |
| chr6 | 29875102 | rs3115619 | - |
| chr14 | 58672348 | rs311845 | + |
| chr10 | 113645856 | rs3127106 | + |
| chr6 | 29872871 | rs3132714 | - |
| chr6 | 29868800 | rs3132718 | - |
| chr12 | 56725452 | rs3214051 | - |
| chr12 | 62763403 | rs342148 | + |
| chr10 | 69324130 | rs34348007 | + |
| chr18 | 38898108 | rs34434760 | + |
| chr13 | 42214119 | rs347418 | + |
| chr17 | 7365448 | rs34755955 | + |
| chr8 | 65191690 | rs34994298 | + |
| chr20 | 17927493 | rs35016526 | + |
| chr1 | 212276874 | rs351417 | - |
| chr20 | 60341506 | rs354741 | + |
| chr7 | 65068922 | rs35794351 | + |
| chr12 | 112630967 | rs36030379 | + |
| chr3 | 114236317 | rs3732782 | - |
| chr1 | 117100373 | rs3754104 | - |
| chr17 | 68206938 | rs3760274 | - |
| chr20 | 3795528 | rs3761218 | + |
| chr1 | 8974362 | rs3765964 | - |
| chr1 | 8951527 | rs3765968 | - |
| chr1 | 201509148 | rs3767525 | - |
| chr15 | 101391959 | rs3784481 | - |
| chr20 | 17466709 | rs3790338 | - |
| chr14 | 44506307 | rs3809430 | - |
| chr1 | 17071812 | rs3818048 | - |
| chr7 | 15873963 | rs38205 | + |
| chr4 | 902698 | rs3822023 | - |
| chr5 | 41573733 | rs3863154 | - |
| chr5 | 103674293 | rs3909338 | - |
| chr1 | 36480052 | rs3917924 | - |
| chr6 | 98785572 | rs3923049 | + |
| chr3 | 192230660 | rs3925222 | - |
| chr9 | 130409635 | rs3926856 | - |
| chr5 | 31682580 | rs39791 | + |
| chr1 | 62221432 | rs4019803 | - |
| chr20 | 45751708 | rs405247 | - |
| chr6 | 24803597 | rs4074236 | - |

| | | | |
|---|---|---|---|
| chr22 | 24010332 | rs422674 | + |
| chr3 | 124242241 | rs4234217 | + |
| chr11 | 72571781 | rs426117 | - |
| chr17 | 3036288 | rs4264423 | + |
| chr5 | 107058155 | rs4269278 | + |
| chr4 | 132737074 | rs4373171 | + |
| chr2 | 136689878 | rs4419275 | + |
| chr11 | 61549915 | rs441938 | - |
| chr12 | 12879853 | rs4462437 | + |
| chr10 | 34905460 | rs4469833 | + |
| chr6 | 74073586 | rs4495244 | + |
| chr9 | 8225531 | rs4497021 | + |
| chr7 | 22112350 | rs4598147 | + |
| chr16 | 82799344 | rs4598906 | + |
| chr1 | 182140796 | rs4652689 | + |
| chr2 | 19791948 | rs4666348 | + |
| chr6 | 31195186 | rs4713447 | + |
| chr7 | 64991278 | rs4718177 | + |
| chr7 | 23789469 | rs4722272 | + |
| chr9 | 69009361 | rs4745500 | + |
| chr12 | 130117883 | rs4760065 | + |
| chr6 | 163118257 | rs476653 | + |
| chr13 | 106406156 | rs4772781 | + |
| chr15 | 93382098 | rs4777857 | + |
| chr17 | 77595906 | rs4789469 | + |
| chr17 | 55304435 | rs4793788 | + |
| chr20 | 24633595 | rs4813525 | + |
| chr2 | 18824642 | rs4832456 | + |
| chr1 | 151809931 | rs4845587 | + |
| chr2 | 69115401 | rs4854547 | + |
| chr9 | 86657853 | rs4878000 | + |
| chr16 | 77701577 | rs4887918 | + |
| chr14 | 101174641 | rs4900487 | + |
| chr6 | 11621871 | rs490280 | + |
| chr15 | 25558943 | rs4906971 | + |
| chr1 | 18852610 | rs4920564 | + |
| chr2 | 44096960 | rs4952699 | + |
| chr3 | 187845171 | rs503787 | + |
| chr1 | 115494175 | rs506049 | + |
| chr6 | 106369142 | rs510405 | - |
| chr18 | 77250689 | rs5374 | + |
| chr4 | 9800648 | rs56139970 | + |
| chr10 | 101280781 | rs56171414 | + |
| chr16 | 88281952 | rs56302546 | + |
| chr3 | 9556313 | rs56407147 | + |
| chr2 | 54692088 | rs56898160 | + |
| chr22 | 37071352 | rs5756506 | + |
| chr11 | 76172117 | rs57946456 | + |
| chr12 | 10733168 | rs5796416 | + |
| chr13 | 36928460 | rs582091 | - |
| chr3 | 3125400 | rs59286216 | + |
| chr4 | 47999036 | rs59363169 | + |
| chr22 | 45051049 | rs6007434 | + |
| chr3 | 8760830 | rs60345038 | + |
| chr20 | 88453 | rs6039403 | + |
| chr20 | 2558741 | rs6050063 | + |
| chr20 | 35006423 | rs6060151 | + |
| chr20 | 32004423 | rs6061087 | + |
| chr20 | 48149399 | rs6066644 | + |
| chr20 | 42344620 | rs6102788 | + |
| chr1 | 199407986 | rs612795 | + |
| chr8 | 86092687 | rs61629815 | + |
| chr15 | 26976494 | rs61999630 | + |
| chr21 | 31590487 | rs62222920 | + |
| chr4 | 65731761 | rs62299607 | + |
| chr4 | 175232180 | rs62337024 | + |
| chr7 | 143418968 | rs62472729 | + |
| chr7 | 143432213 | rs62474772 | + |
| chr1 | 233618245 | rs635698 | - |
| chr11 | 117293108 | rs638405 | + |
| chr11 | 78153964 | rs643430 | + |
| chr3 | 114236473 | rs6438191 | + |

| | | | |
|---|---|---|---|
| chr3 | 116015687 | rs6438296 | + |
| chr3 | 125009782 | rs6438869 | + |
| chr3 | 45124731 | rs6441891 | + |
| chr8 | 33498556 | rs6468171 | + |
| chr8 | 65178463 | rs6472179 | + |
| chr10 | 18902718 | rs6481583 | + |
| chr15 | 66957020 | rs6494621 | + |
| chr16 | 27813510 | rs6498038 | + |
| chr12 | 120764559 | rs6553 | + |
| chr5 | 165709385 | rs6556822 | + |
| chr12 | 120878449 | rs656933 | - |
| chr14 | 80457125 | rs6574576 | + |
| chr11 | 100069001 | rs6590574 | + |
| chr1 | 4459129 | rs662036 | - |
| chr2 | 39162037 | rs66771682 | + |
| chr1 | 159883032 | rs6682796 | + |
| chr4 | 115302551 | rs66997216 | + |
| chr6 | 116250092 | rs6755 | - |
| chr3 | 54601810 | rs6765694 | + |
| chr3 | 39847010 | rs6777708 | + |
| chr3 | 157407795 | rs6803628 | + |
| chr4 | 119025980 | rs6828669 | + |
| chr4 | 179766847 | rs6844851 | + |
| chr5 | 178538695 | rs688074 | + |
| chr6 | 125316311 | rs6899690 | + |
| chr6 | 1137899 | rs6904803 | + |
| chr6 | 36290890 | rs6914153 | + |
| chr6 | 131145967 | rs6942184 | + |
| chr7 | 18193559 | rs6945251 | + |
| chr7 | 15629375 | rs6958841 | + |
| chr12 | 130073404 | rs697792 | + |
| chr9 | 78927489 | rs7030136 | + |

| | | | |
|---|---|---|---|
| chr9 | 90808223 | rs7036417 | + |
| chr9 | 11039639 | rs7044282 | + |
| chr10 | 3616727 | rs705469 | + |
| chr10 | 129833518 | rs7069546 | + |
| chr2 | 154370165 | rs707040 | + |
| chr10 | 126163546 | rs7077301 | + |
| chr10 | 12547151 | rs7078960 | + |
| chr11 | 48489102 | rs7103557 | + |
| chr8 | 17313520 | rs71212685 | - |
| chr11 | 12992289 | rs7129181 | + |
| chr11 | 111185891 | rs7130452 | + |
| chr15 | 78841397 | rs7166501 | + |
| chr15 | 33213807 | rs7169713 | + |
| chr16 | 12442437 | rs7185322 | + |
| chr16 | 72090258 | rs7190995 | + |
| chr16 | 79115745 | rs7191506 | + |
| chr16 | 78590110 | rs7200682 | + |
| chr16 | 2013618 | rs7200747 | + |
| chr17 | 73535958 | rs7209879 | + |
| chr17 | 67029895 | rs7216886 | + |
| chr18 | 7067653 | rs7234111 | + |
| chr18 | 78440167 | rs7241158 | + |
| chr14 | 89727083 | rs72689472 | + |
| chr1 | 202264291 | rs727724 | + |
| chr6 | 155582297 | rs72994676 | + |
| chr12 | 63621661 | rs7311307 | + |
| chr13 | 39346149 | rs7318557 | + |
| chr13 | 74345518 | rs7333414 | + |
| chr13 | 103977322 | rs7338623 | + |
| chr16 | 29230794 | rs735645 | - |
| chr7 | 10888515 | rs7357330 | + |
| chr8 | 70524140 | rs7387249 | + |

| | | | |
|---|---|---|---|
| chr11 | 48497341 | rs7395662 | + |
| chr6 | 9413276 | rs742511 | - |
| chr1 | 25875935 | rs74378533 | + |
| chr5 | 175331195 | rs7445817 | + |
| chr2 | 225173107 | rs751264 | + |
| chr1 | 230452281 | rs7514115 | + |
| chr1 | 247599026 | rs7523061 | + |
| chr1 | 49017536 | rs7524637 | + |
| chr1 | 21514216 | rs7535497 | + |
| chr1 | 47247503 | rs7552123 | + |
| chr2 | 238927379 | rs7571467 | + |
| chr2 | 64538340 | rs7581658 | + |
| chr17 | 34606024 | rs758409 | - |
| chr2 | 46746261 | rs7584870 | + |
| chr3 | 77472155 | rs7614439 | + |
| chr20 | 23446011 | rs761724 | - |
| chr3 | 7469204 | rs7623055 | + |
| chr3 | 101546979 | rs7628681 | + |
| chr3 | 60116459 | rs7648665 | + |
| chr6 | 51748379 | rs765525 | - |
| chr2 | 73151113 | rs768851 | + |
| chr4 | 41798742 | rs7689428 | + |
| chr4 | 4689125 | rs7692353 | + |
| chr5 | 144145201 | rs7716084 | + |
| chr6 | 161778667 | rs7754162 | + |
| chr8 | 31898373 | rs776378 | + |
| chr6 | 69755811 | rs7775172 | + |
| chr7 | 133120296 | rs7777197 | + |
| chr7 | 138632991 | rs7792330 | + |
| chr7 | 12551849 | rs7800736 | + |
| chr8 | 139979861 | rs7837321 | + |
| chr9 | 1170227 | rs7848270 | + |

FIGURE S1

GENOME SCANS OF Z-SCORES, MFED DIFFERENCES AND SEQUENCE DIVERGENCE

OF HCV GENOTYPES 1a, 1b AND 3a

A)

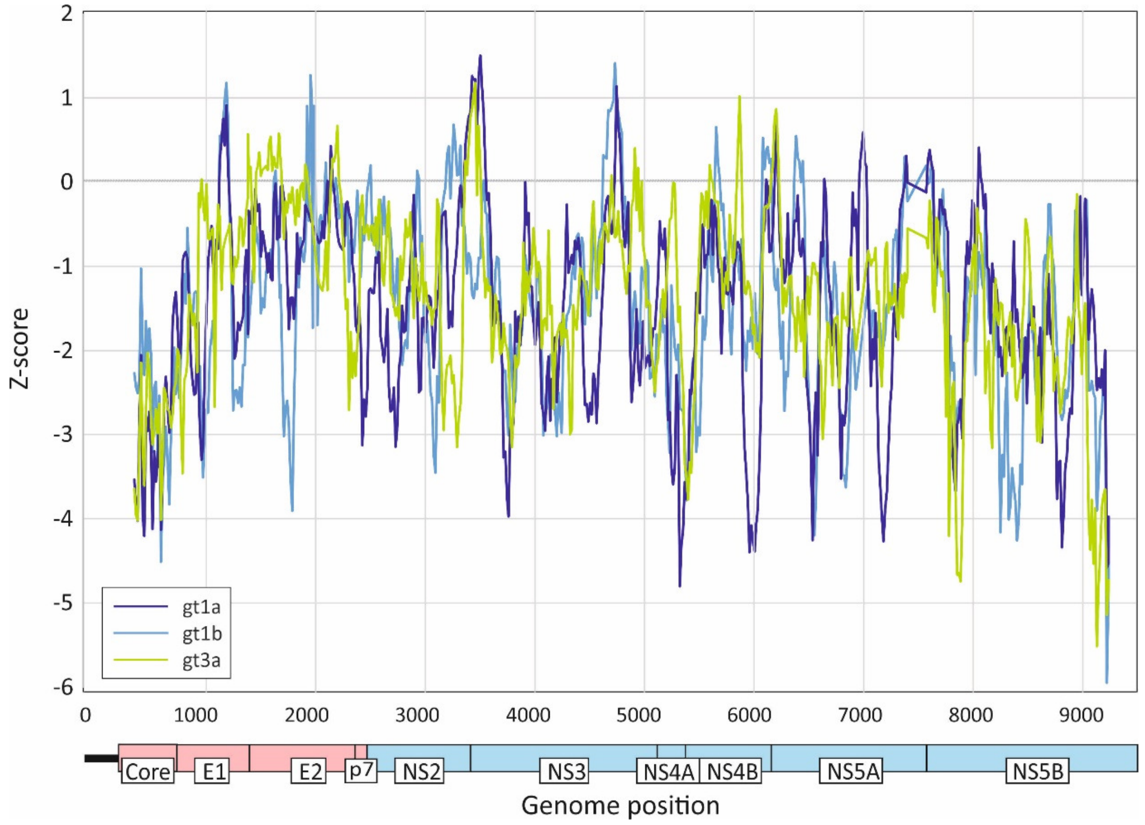


B)

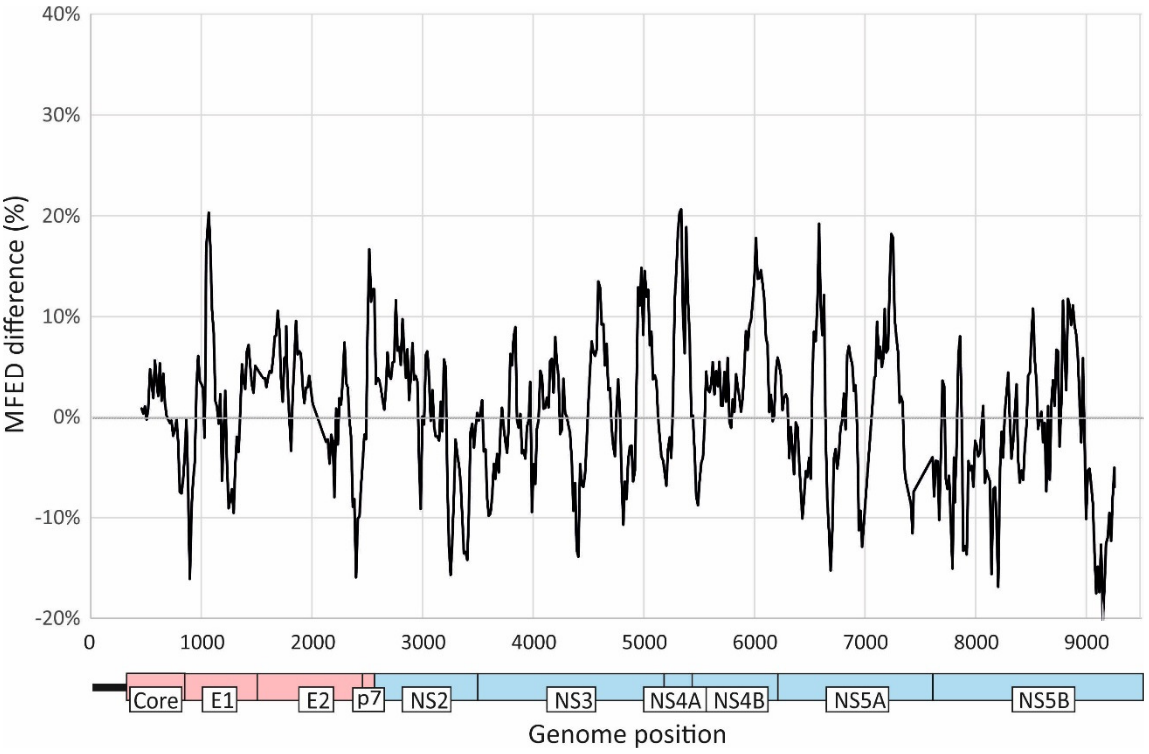

C)

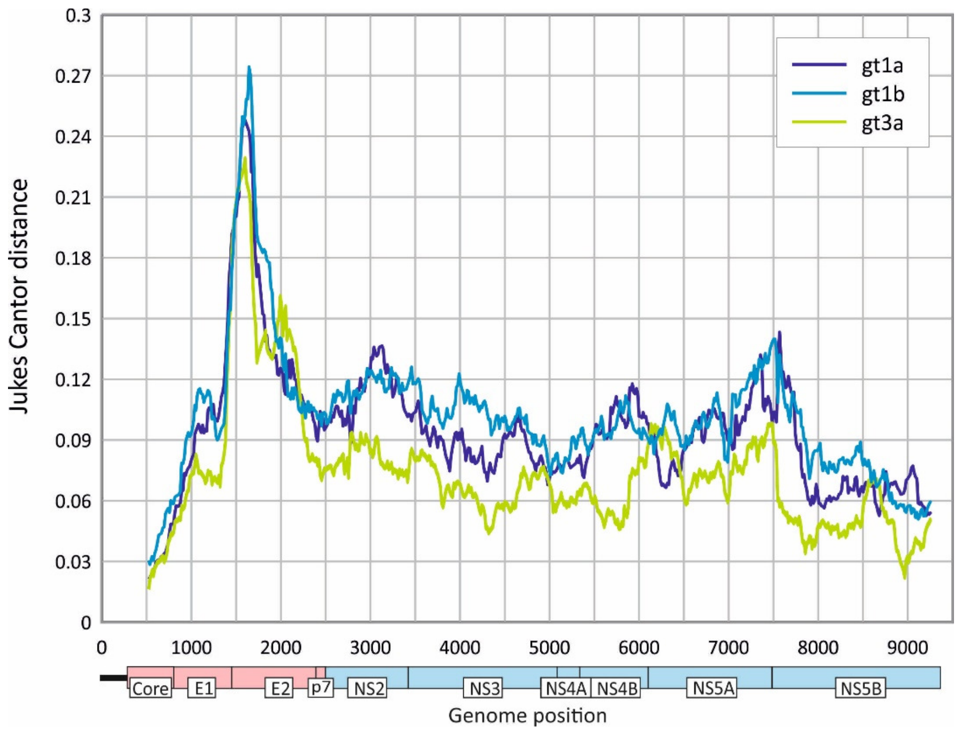


A) Z-scores calculated for consecutive 240 base fragments, incrementing by 15 bases across the HCV coding region (569 fragments). Z-scores represent the position of the MFE of the native sequence within the distribution of MFE values of 49 sequence order randomised controls. Graph lines show mean values of genotypes 1a (n=388), 1b (n=106) and 3a (n=855) sequences plotted by genome position. (B) Differences in MFED values for gt1a and gt3a scanned across the genome. (C) Mean divergence between prototype sequences of genotype 1a (M67463 – H77), 1b (D90208 – HCV-J) and 3a (D17763 – NZL-1) and representative sequences of each genotype sampled from the study population. Mean distances were calculated for 360 fragments incrementing by 15 bases between fragments with Jukes-Cantor correction for multiple substitutions. .

FIGURE S2

INFLUENCE OF SEQUENCE LENGTHS and SUBOOPTIMAL SAMPLING METHOD ON CONTOUR PLOT PREDICTIONS

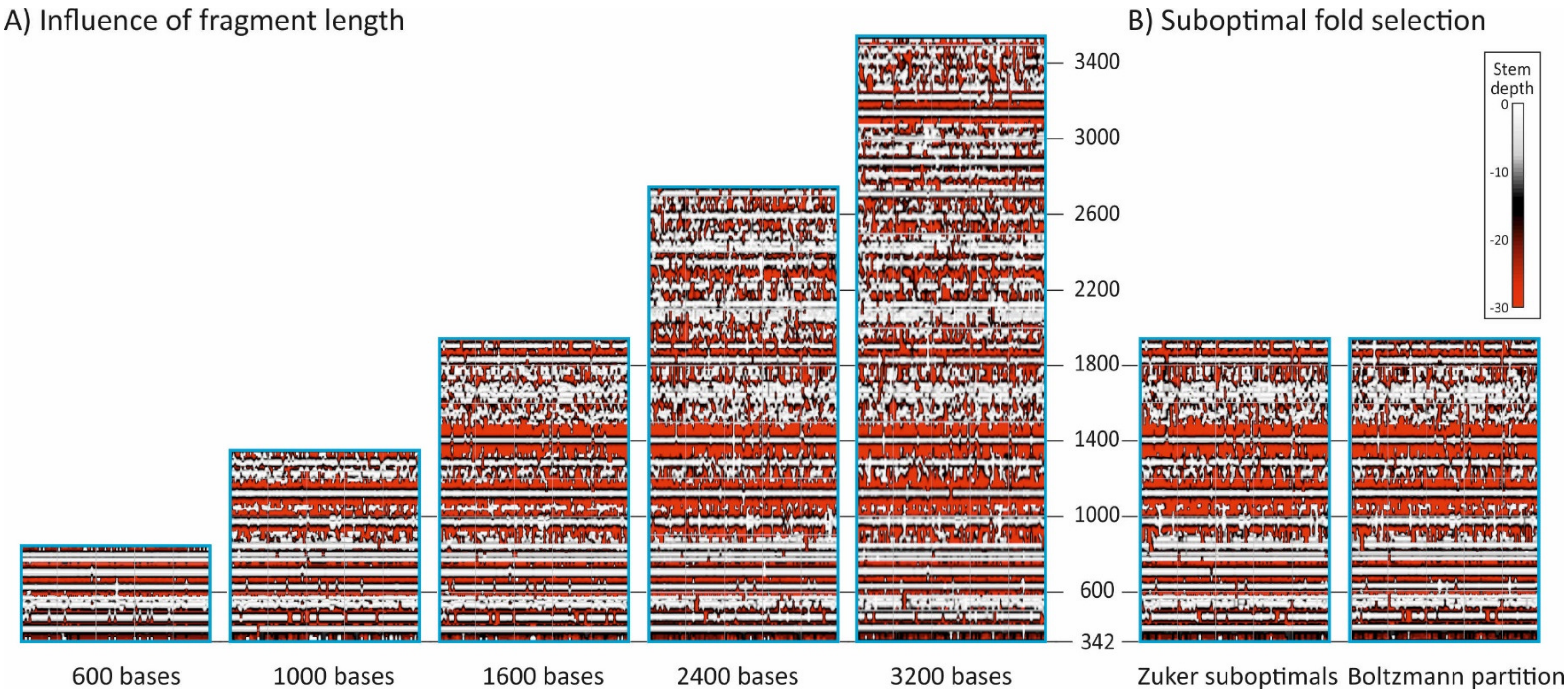


Contour plots of 99 HCV genotype 1b sequences generated using different run parameters. (A) Comparison of contour plots for sequence fragments of lengths 600, 1000, 1600 (as used for other plots presented in the study), 2400 and 3200 bases (approaching the upper bound of sequence length for RNAsubopt). (B) Comparison of the default selection methods for suboptimal folds (those within 5% of the optimum (minimum) MFE (selection of --deltaEnergy parameter in RNAsubopt) sd used in the plots shown in Fig. S2A) with alternative methods based on a random selection of Boltzman weights in the partition function (--stochBT) and on Zuker sub-optimals (--zuker).

FIG. S3. CONTOUR PLOTS OF PROTOTYPE SEQUENCES OF EACH DIFFERENT GENOTYPE AND SUBTPYE OF HCV

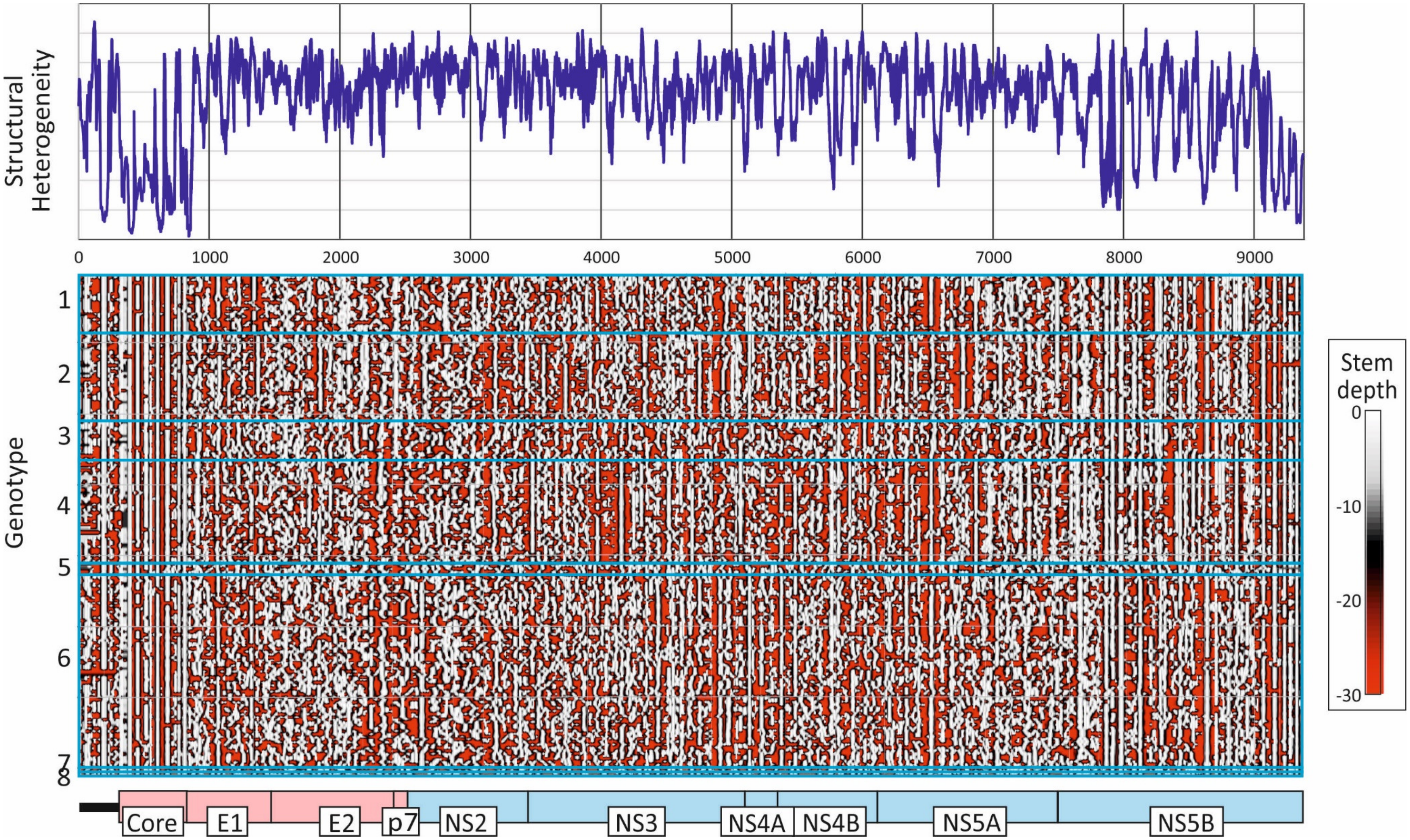


Contour plot of single examples of each currently classified and candidate HCV subtype and genotype (data derived from https://talk.ictvonline.org/ictv:wikis/flaviviridae/w/sg:_flavi/56/hcv-classification) listed in Table S1; Suppl./ Data.

FIGURE S4

MFED VALUES FOR REPRESENTATIVE EXAMPLES OF EACH HCV GENOTYPE / SUBTYPE

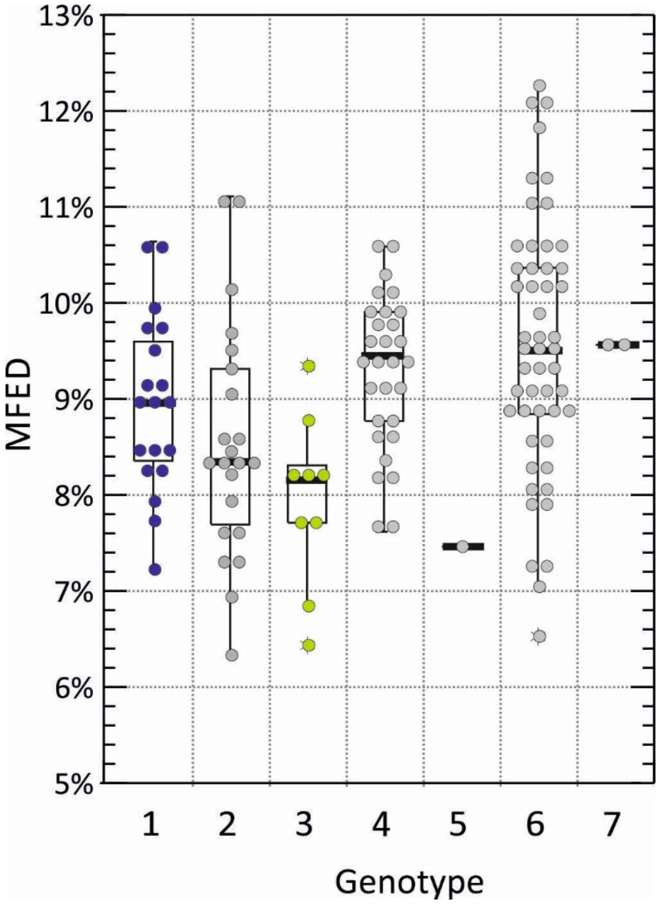


Mean values of sequence fragments for individual polyprotein sequences of all classified subtypes for HCV genotypes 1-7 (individual representative sequences for each subtype shown. The box plots show (from the top): 2 standard deviations (SDs) above mean, 1 SD above mean, mean, 1 SD below mean and 2 SDs below mean; stars represent outliers outside this range.

FIG. S5. CONTOUR PLOTS OF DIFFERENT FMDV TYPES

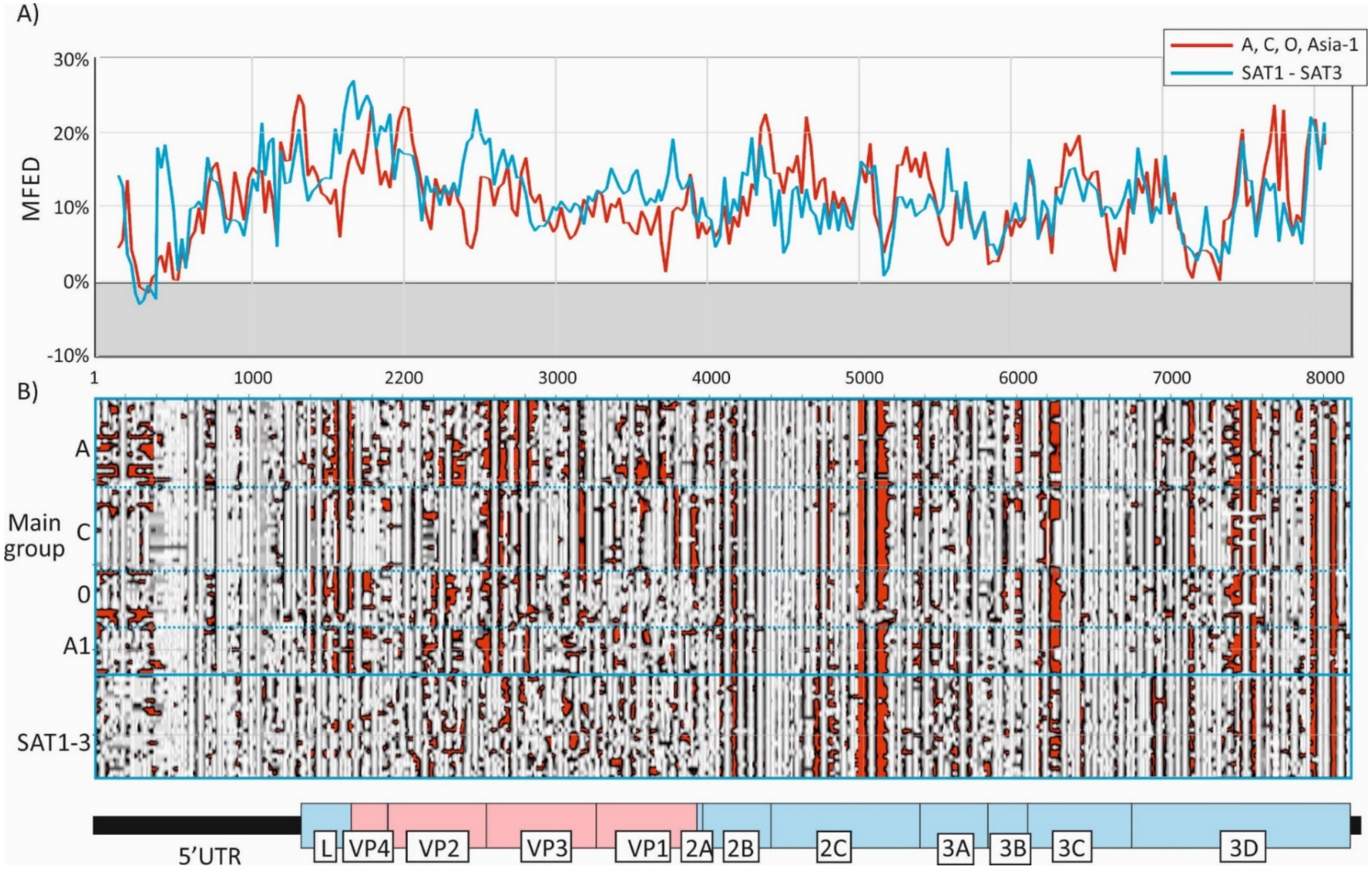


RNA structure prediction in different serotypes of FMDV. (A) MFED scan of the genome of the two FMDV groups using fragment sizes and increments as used for HCV (Fig. 1). (B) Contour plot of representative example sequences of each FMDV type.
